# A Continuous Time Representation of smFRET for the Extraction of Rapid Kinetics

**DOI:** 10.1101/2020.08.28.267468

**Authors:** Zeliha Kilic, Ioannis Sgouralis, Wooseok Heo, Kunihiko Ishii, Tahei Tahara, Steve Pressé

## Abstract

Our goal is to learn kinetic rates from single molecule FRET (smFRET) data even if these exceed the data acquisition rate. To achieve this, we develop a variant of our recently proposed *hidden Markov jump process* (HMJP) with which we learn transition kinetics from parallel measurements in donor and acceptor channels. Our HMJP generalizes the hidden Markov model (HMM) paradigm in two critical ways: (1) it deals with physical smFRET systems as they switch between conformational states in *continuous time*; (2) it estimates the transition rates between conformational states directly without having recourse to transition probabilities or assuming slow dynamics (as is necessary of the HMM). Our continuous time treatment learns transition kinetics and photon emission rates for dynamical regimes inaccessible to the HMM which treats system kinetics in discrete time. We validate the robustness of our framework on simulated data and demonstrate its performance on experimental data from FRET labeled Holliday junctions.

## 1 Introduction

Fluorescence experiments based on single molecule Förster resonance energy transfer (smFRET) can probe the switching kinetics between conformational states defined by different inter- and intra-molecular distances (1–10). In a prototypical intra-molecular smFRET experiment, one portion of a molecule of interest is attached to a *donor* fluorophore and another to an *acceptor* fluorophore (1–18). In such an experiment, the excitation wavelength is most commonly adjusted to excite the *donor* (9, 14, 16–18). For a donor sufficiently far from the acceptor, the donor is excited and emits shorter wavelength light as compared to the longer wavelength light emitted by the acceptor in the case of energy transfer when donor and acceptor are in proximity (9, 14, 16–18). Due to the difference in the wavelength of light emitted by the donor and acceptor, the photons emitted are registered across different detectors (9, 14, 16–18). We refer to the recordings in the two detectors as the donor and acceptor channels (9, 14, 16–18). As such, the sequence of photon detections (2) encodes the kinetics according to which the distance varies between both fluorophores down to the *μ*s timescale (1–9). Existing approaches to analyze smFRET data include FRET efficiency histogram-based methods though such methods discard the important temporal information encoded in donor and acceptor photon arrival sequences (5–9, 11–13, 15, 16, 19). What is more, FRET efficiency histogram analysis is difficult to generalize beyond 2 conformational states as discussed in (20). In addition, on account of the arbitrary binning required to construct histograms, such methods may lead to inconclusive or erroneous estimates (5, 6, 21–27). For these reasons, modeling efforts have moved toward more direct time series analysis (7, 20, 28–31).

By relying on the hidden Markov model (HMM) paradigm (1, 3,7, 10, 30, 32), time series analysis fully considers the temporal arrangement (i.e., the sequence) of photon detections and avoids histogram binning artifacts (1, 3,7, 10, 32). This paradigm is especially fruitful in analyzing demanding experiments as the conformational states, treated as the hidden states of the HMMs, are themselves indirectly observed due to shot-noise (1, 3) and also allows for inclusion of measurement noise and specialized detector characteristics in the analysis (1, 3,7, 10, 32).

HMMs and their variants, starting from (15) and motivating latter efforts such as the H^2^MMs (1, 3,30, 35, 37) and bl-ICON (7), are the state-of-the-art in the analysis of smFRET data. However, HMMs have built into them important assumptions that we wish to lift in an effort to analyze conformational transitions occurring on time scales faster than the data acquisition rate 1/Δ*t* supported by the detectors. Before describing these assumptions, we discuss Δ*t* (the bin size for photon collection) otherwise known as frame rate or temporal resolution. As Δ*t* is determined mostly by the detectors, for simplicity we assume that the same Δ*t* applies on both channels.

In order to learn about rates exceeding the data acquisition rate, the most critical HMM assumptions that we need to lift are the following:

- HMMs make the assumption that the state switches occur *rarely* as compared to Δ*t*. In other words, all switching rates are assumed much slower than 1/Δ*t*. For this reason, HMMs are formulated using transition probabilities rather than transition rates.
- HMMs make the assumption that transitions occur *precisely* at the end of each data acquisition period (30, 35, 38–43). In other words, intra-frame motion does not occur. For this reason, HMMs represent only instantaneous states and measurements.

The former assumption is particularly relevant to the present study as it requires, prior to the analysis, that all transitions are slower than Δ*t*. In general, this is not only unknown but also practically difficult to quantify beforehand. After all, the objective of many experiments is to determine the switching rates in the first place. The latter assumption is also relevant to the present study as, prior to the analysis, it needs to balance two competing requirements: an upper bound on Δ*t* set by the fastest switching rate and a lower bound on Δ*t* supported by the detection hardware.

There have been few theoretical approaches proposed to overcome these challenges. For example, in the H^2^MM (3), a time grid finer than the measurement time interval is imposed and a HMM is implemented on this finer grid (3). This method is known to suffer from computational complexity (44) and it is for this reason that formulations have been sought in continuous time (44, 45). Motivated by the H^2^MM (3), our goal is to move to an exact continuous time treatment for smFRET systems with no approximation on the timescales of the switching kinetics.

In Fig. 1 panels (a1),(b1), we illustrate an example of smFRET measurements. In panel (a1), we show an example with slow kinetics (relative to Δ*t*). These data are primarily contaminated with shot-noise and can be reliably analyzed within the HMM paradigm. On the other hand, in panel (b1), we demonstrate an example with fast kinetics (relative to Δ*t*). On top of shot-noise, these data are contaminated with significant intra-frame transitions, consequently they cannot be analyzed within the HMM paradigm. As the molecule undergoes fast switching between the states during a data acquisition period, the measurement associated with that period reports on the average signal from the molecule over that period and the state of the molecule cannot be deduced from a model operating exclusively on instantaneous states such as within the HMMs paradigm. Nonetheless, the information about the fast switching is itself encoded within that measurement. It is therefore necessary to develop a method that can accommodate *continuously* evolving dynamics in order to learn the kinetics from the measurements instead of invoking an HMM formulation that, by construction, violates this critical feature.

**Fig. 1.**
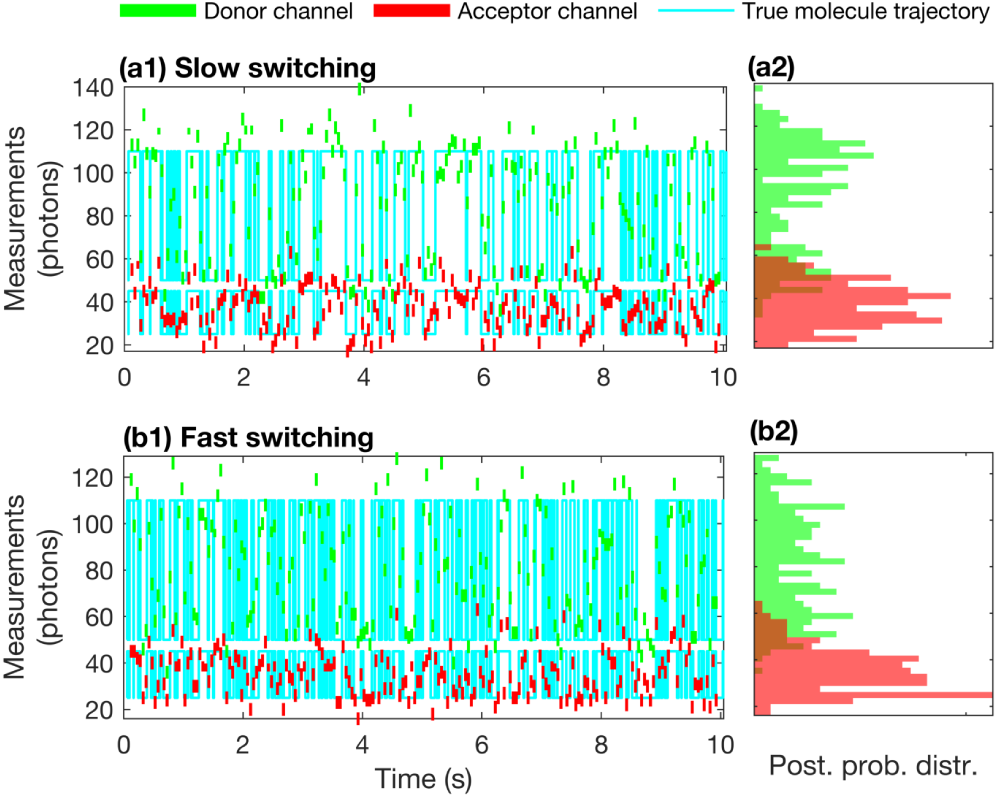
An illustration of single molecule switching kinetics and corresponding measurements. In panels (a1) and (a2), we provide the simulated trajectory (cyan) of the single molecule between two states (σ_1_, σ_2_). The photon emission rates, i.e., number of emitted photons per unit time in the absence of noise in both donor and acceptor channels associated with the conformation states σ_1_ and σ_2_ are labeled with 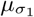 and 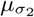, respectively. This simulated experiment provided in this figure starts at t_0_ = 0.05 s and ends at t_N_ = 10 s with data acquisition period Δt = 0.05 s. Here in panels (a1) and (b1), for visual purposes, we assume that the measurements are acquired by a detector with fixed exposure period τ = 50 ms in donor (green) and acceptor (red) channels. As the molecule switches between states σ_1_, σ_2_ during an integration period, the measurements represent the number of emitted photons that capture the average of the photons emitted with rates 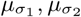 associated with the visited states. In this figure, panels (a1)-(a2) represent the simulated data where the molecule switching kinetics between the conformation states are slower than the data acquisition rate. On the other hand, in panels (b1)-(b2), we demonstrate a single molecule trajectory (cyan) when the molecule’s switching kinetics are faster than the data acquisition rate. In panel (a2), slow kinetics of the molecule give rise to well separated state occupancy histograms in donor/acceptor channels around the average photon emission rates. By contrast, in panel (b2), we don’t observe well separated histograms due to fast switching kinetics of the molecule.

To achieve this, a new method needs to: i) represent the dynamics of the molecule in continuous time; ii) model the acquired measurements at each data acquisition times via the average dynamics of the molecule within the associated data acquisition period; and iii) entail manageable computational cost so it allows for practical applications.

For this reason, we now turn to Markov jump processes (MJPs) describing continuous time dynamics (46–50) for which recent developments in computational statistics have suggested strategies for inferring rates using MJPs from traditional, continuous time, data (44, 45, 51). The main challenge presented by smFRET experiments, however, lie in the fact that measurements do not directly report back on the fast kinetics of molecules. Rather, frame rates report on the *average* state of the molecule over the Δ*t* exposure window. This problem is exacerbated for molecular kinetics fast as compared to Δ*t*. The focus of this work is really to validate a workaround, that we call the Hidden MJP (HMJP), to this critical challenge. As we will see, the HMJP is a generalization of the HMM. As such, the HMJP mathematically exactly reduces to the HMM in the limit that exposure period Δ*t* → 0.

To achieve this, in Section 2, we start with our HMJP smFRET model description and also, briefly for sake of comparison, summarize plain HMMs for smFRET applications. Next, in Section 3, we provide HMM and HMJP analysis comparisons for simulated measurements. In the comparison of these two methods, we present their performances in learning photon emission rates, transition probabilities (for HMMs) and kinetic rates (for HMJPs). We demonstrate how HMJPs successfully outperform HMMs especially for fast kinetic rates as compared to the data acquisition rate. Their comparison on slow switching kinetics of simulated smFRET data (where both HMM and HMJP expectedly do well) is relegated to Appendix A. Subsequently, we move onto the analysis of experimental data. We provide a comparison of HMJPs and HMMs in learning molecular trajectories, transition probabilities and kinetic rates. Lastly, in Section 4, we discuss the broader potential of HMJPs for smFRET applications.

## 2 Methods

Below, we provide the mathematical description of a physical system that models smFRET experiments. Subsequently we use this model in conjunction with given data to extract our estimates.

### 2.1 Model Description

#### 2.1.1 Dynamics of the System

We denote the trajectory, describing the molecule’s conformational states at any given time, with 𝒯(·). Here, 𝒯 (*t*) corresponds to the molecule’s state at time *t*. Thus 𝒯 (*t*) is a *function* over the time interval [*t*_0_, *t*_*N*_]. The accessible states of the molecule are labeled with *σ*_*k*_ and indexed with *k* = 1, …, *K*. For example, these states can be considered the *K* = 2 isomerization states of Holliday junctions (3). If the molecule is in state *σ*_*k*_ at time *t*, then we denote it by 𝒯 (*t*) = *σ*_*k*_, (44, 45).

We model the transitions of the molecule as a *memoryless* process (35, 50, 52, 53) and assume exponentially distributed waiting times in each state. Together, these define our *Markov process*. In particular, as we model waiting times in continuous times, our Markov process is a *Markov jump process* (50).

In greater detail, we assume that the system chooses stochastically a state *σ*_*k*_ at the onset of the experiment that is denoted by 𝒯 (*t*_0_) = *σ*_*k*_. The probability of determining this initial state *σ*_*k*_ is labeled with 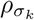. The collection of all initial probabilities is a probability vector and labeled with 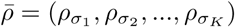, (54–57).

Switching rates fully describe the switching kinetics of the molecule and they are labeled with 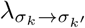 for all possible states *σ*_*k*_,*σ*_*k*′_ where *k, k*^′^ = 1, 2, …, *K* and 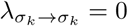 by definition for *k* = 1, 2, …, *K*. For mathematical convenience, we keep track of the *escape* rates as an alternative parametrization of the switching kinetics which are given by

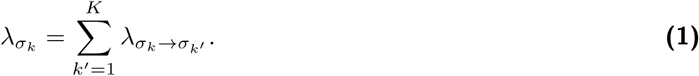

We collect all escape rates in 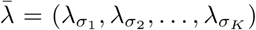. Moreover, for computational convenience, we track the normalized switching rates by the escape rates

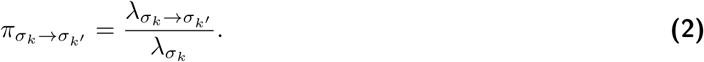

The collection of all normalized switching rates from state *σ*_*k*_ is denoted by 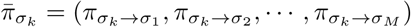. We see that each 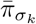 forms a probability vector (56). We can gather all transition probabilities in a matrix 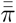 that reads

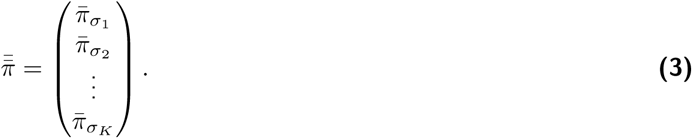

Given 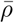 and 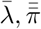, the trajectory 𝒯 (·) is obtained by a variant of the Gillespie algorithm (52) which determines a succession of states for the conformations of the system *s*_0_, *s*_1_, · · ·, *s*_*M*−1_ and their durations *d*_0_, *d*_1_, *d*_2_, · · ·, *d*_*M*−1_. These together define 𝒯 (·) throughout the time course [*t*_0_, *t*_*N*_] by

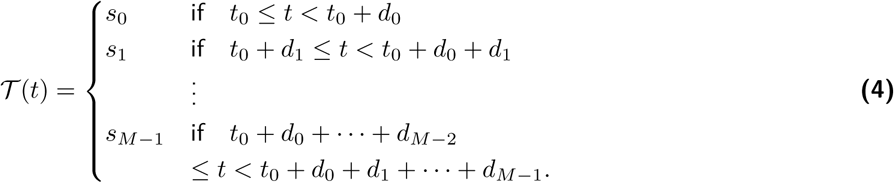

For clarity, we encode 𝒯 (·) in a triplet 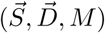, where 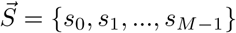 and 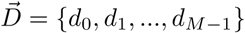 and *M* is the size of 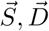.

#### 2.1.2 Measurements

The measurements in a typical smFRET experiment report on the conformational state of the molecule as it changes through time. These come in the form of two time series: 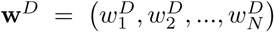 and 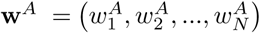 which are the recordings in the donor and acceptor channels, respectively. In particular, the subscripts here indicate the time level of measurements. For clarity, we assume that the measurements are time-ordered, so the *n* = 1 label coincides with the earliest acquired measurement and the *n* = *N* label coincides with the latest.

We assume that measurements occur on a regular time interval denoted by Δ*t* (although the switching of the molecule between states occurs in continuous time). Here, the values of 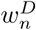 and 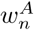 report on a number of photons collected between *t*_*n*− 1_ and *t*_*n*_ where *t*_*n*_ = *t*_*n*− 1_ + Δ*t*. For completeness, we introduce an additional time level *t*_0_ which precedes the first measurement time *t*_1_. This time level, *t*_0_ defines the experiment’s onset, which is not associated to any measurement; see Fig. 1.

One of the common assumptions of HMMs is that the instantaneous state of the molecule at *t*_*n*_ determines the measurements 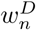 and 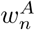. However, for realistic detectors, the reported values 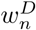 and 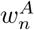 are affected by the entire photon trajectory of the molecule during the *n*^th^ integration period represented by the time window [*t*_*n*_ − *τ, t*_*n*_]. Here, *τ* is the duration of each integration time for fluorescence experiments.

When we supplement our dynamical model (fully described in Section 2.1.1) with measurements, we must include a distribution describing the measurement statistics. We do so by first discussing the state specific photon emission rates in the donor and acceptor channels which we label with 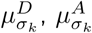, where the subscript highlights the state dependence of the photon emission rate. For simplicity, we gather the state specific photon emission rates in 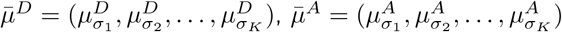.

If the molecule remains in a single state *σ*_*k*_ throughout an entire exposure period [*t*_*n*_ − *τ, t*_*n*_], then the detector is triggered by 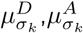 and ambient contributions (background) which we label with 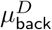 and 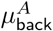 for the donor and acceptor channels, respectively. As such, the reported measurement, 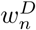, is similar to 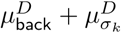 and 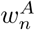 is similar to 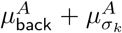. However, if the molecule switches between multiple states during the same exposure period, the detector is influenced by the levels of every state attained. More specifically, the *n*^th^ binned photon counts triggering the detector during the *n*^th^ exposure period, [*t*_*n*_ − *τ, t*_*n*_], is obtained from the 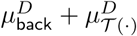 in the donor channel and 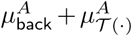 in the acceptor channel over this exposure. Mathematically, this is equivalent to 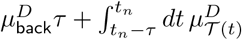 in the donor channel and 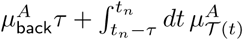 in the acceptor channel.

With measurement noise, such as shot-noise (22, 58–61), quantification noise (62–64), or amplification noise in the case of EMCCD detectors (7, 65–67) and other degrading effects that are common in the detectors currently available, each measurement 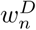 and 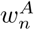 depends *probabilistically* upon the triggering signal (68–70). Of course, the precise relationship depends on the detector employed in the experiment and differs between the various types of cameras, single photon detectors or other devices used. Here, we continue with a shot-limited formulation, which results in

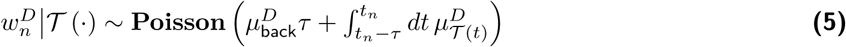

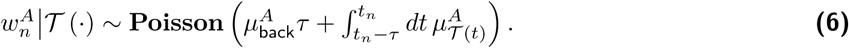

#### 2.1.3 FRET Efficiency

Later we will be making use of the notion of FRET efficiency. For this reason we define two different types of FRET efficiencies: the *characteristic* FRET efficiency (7, 15, 71–75) as well as the *apparent* FRET efficiency (7, 15, 71–75). Characteristic FRET efficiency (7, 15, 71–75) labeled with 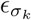 for *k* = 1, 2, …, *K* is defined as follows

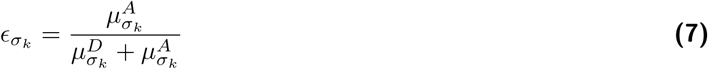

and only depends on the conformational states (7, 15, 71–75). By contrast, the apparent FRET efficiency (which is often used as a proxy for 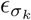) is defined as follows

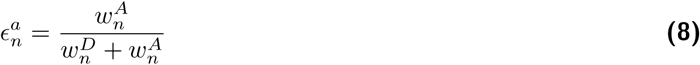

for *n* = 1, 2, …, *N*. Unlike the former, the latter is affected by artifacts such as measurement noise and background (15). As a result, the apparent efficiency may attain different values over time even when the molecule does not switch its conformational state.

### 2.2 Model Inference

Our goal is to learn the initial probabilities 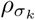, photon emission rates 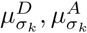, switching rates 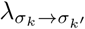 for all states and the trajectory of the system 𝒯 (·) during the full time course [*t*_0_, *t*_*N*_] of the experiment by using the measurements **w**^*D*^, **w**^*A*^, and the model associated with the smFRET experiment that has just been described. First, we will explain how we learn these quantities using time series analysis within the naive HMM paradigm (7, 74, 75). Subsequently, we will present how we tackle smFRET data using our continuous time HMJPs.

#### 2.2.1 Model Inference via HMMs

HMMs inherently assume that each measurement 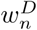 and 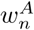 acquired at time *t*_*n*_ depends only on the molecule’s state at the time of data acquisition namely 𝒯 (*t*_*n*_). Therefore, we have the following approximations for the HMM formulation 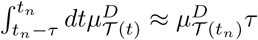 and 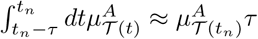 during the exposure period [*t*_*n*_ − *τ, t*_*n*_]. Consequently, Eq. (5) and Eq. (5) become

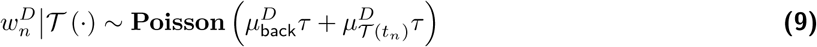

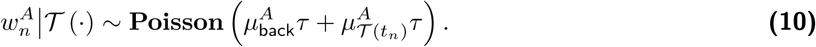

Given the HMM formulation and the smFRET data **w**^*D*^, **w**^*A*^, we can *directly* learn the transition probabilities that govern the transitions of the molecule between its conformational states at any data acquisition time *t*_*n*_ for all *n* = 1, 2, …, *N*. We label a molecule’s state within the HMM paradigm at time *t*_*n*_ with 𝒯 (*t*_*n*_) = *c*_*n*_. For clarity, transition probabilities determine the switching probabilities for a molecule’s transitions denoted by *c*_*n*−1_ → *c*_*n*_ → *c*_*n*+1_. Within the HMM formulation, 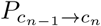 denotes the transition probability of the molecule from state *c*_*n*−1_ to *c*_*n*_.

Given that there are *K* conformational states for the molecule, the number of transition probabilities 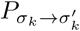 becomes *K*^2^. We gather the transition probability from conformational state *σ*_*k*_ to any other conformational state including itself in 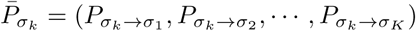 that is normalized as a probability vector (namely each component sums up to 1). Eventually, we gather all transition probability vectors in a matrix called transition probability matrix denoted by 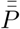 which reads as follows

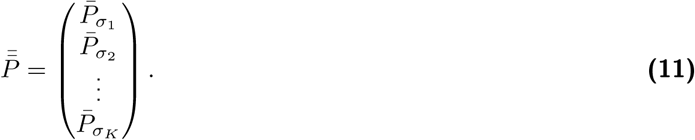

There is a connection between the transition probability matrix 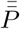 and the molecule’s switching rates 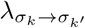 and escape rates 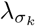. Provided that we gather the switching rates and escape rates in the following matrix, called *generator*,

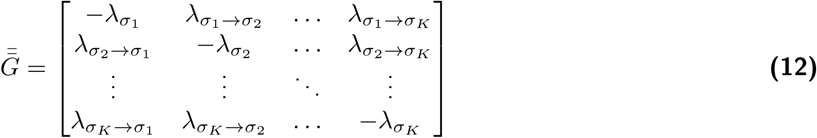

then 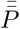 can be calculated based on 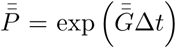 where exp (·) stands for the matrix exponential. Knowing 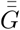 determines 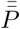 uniquely; however, knowing 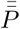 provides only a proxy for 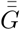. Specifically, if we assume that 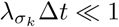 then we can use an approximation

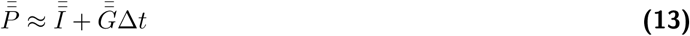

where 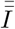 is an identity matrix of size *K* ×*K*. Thus we can calculate an approximation for 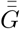 that is 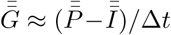.

Now, we provide the formulation for how to estimate the quantities including initial probabilities, 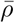, transition probabilities 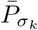, photon emission rates 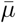 and the trajectory of the system 𝒯 (·) which is encoded by 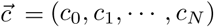, within an HMM paradigm. The HMM formulation is governed by the following statistical model

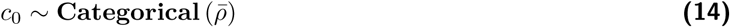

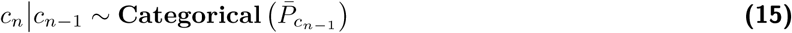

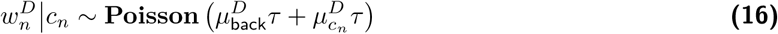

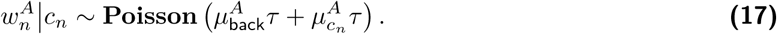

From now on, we follow the *Bayesian paradigm* (54, 76) which requires us to prescribe prior distributions for the parameters.

We start with the prior distributions placed on the transition probabilities 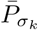 for all *k* = 1, 2, …, *K* and 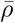. We choose to place Dirichlet distributions with concentration parameters *A* and *a* for 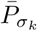 and 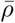, respectively, that are conjugate to the Categorical distribution (30, 35, 57, 77) and formulated as follows

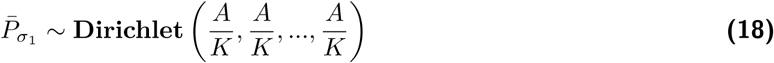

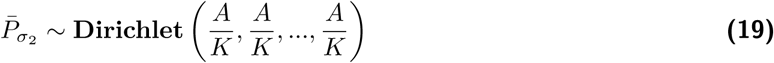

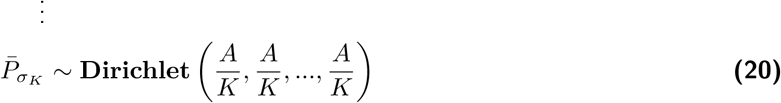

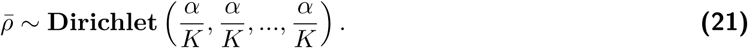

Subsequently, we place priors on the photon emission rates 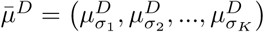 and 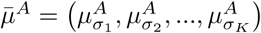. The prior that we choose to place is the Gamma distribution as it has positive support

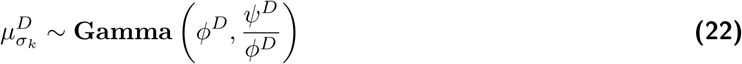

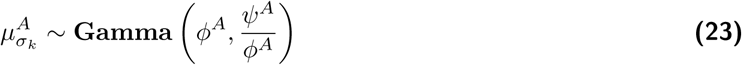

with hyperparameters *φ*^*D*^, *ψ*^*D*^, *φ*^*A*^, *ψ*^*A*^.

We estimate the background photon emission rates 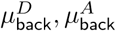 by separate measurements that contain only ambient contributions as we explain in Supporting Material (A). These measurements can be obtained either after both donors and acceptors photobleach or, in a separate experiment, in which no FRET labeled molecule is present.

With all priors specified, we form the full posterior distribution (7, 30, 31, 35, 37, 77, 78)

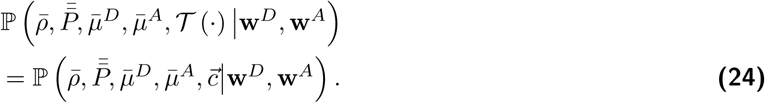

Since we do not have an explicit formula for Eq. (24) we build a custom MCMC sampling scheme to generate pseudorandom numbers from Eq. (24). Details of the computational scheme are provided in Section 2.2.3.

#### 2.2.2 Model Inference via HMJPs

The HMJP does not require any approximations on the kinetics regime allowed and instead applies directly on the formulation provided in Eq. (5) and Eq. (6). Just as with the HMM, for the HMJP formulation, we also operate within the *Bayesian paradigm* (54, 76) and thus place prior distributions on 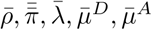.

First, we prescribe the prior distributions on the escape rates 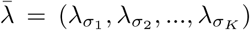. For these, we choose Gamma distributions which are conjugate to the exponentially distributed holding times, i.e.,

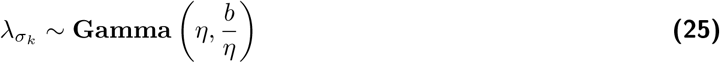

for all *k* = 1, 2, …, *K* with hyperparameters *η, b*. Subsequently, we prescribe independent conjugate Dirichlet distributions on the transition probabilities 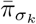 for all *k* = 1, 2, …, *K*

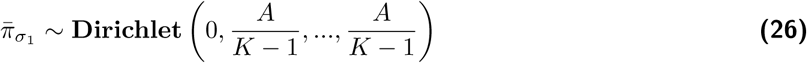

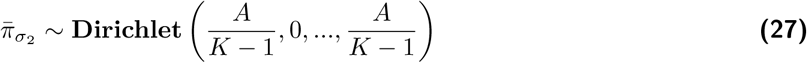

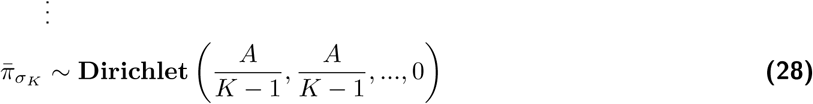

with concentration hyperparameter *A*. Prior distributions placed on 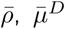 and 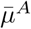 are provided in Eq. (21) and Eq. (23), respectively.

The full posterior distribution for the HMJP formulation is

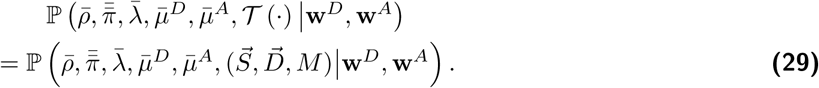

Since we do not have an analytical formula for Eq. (29), we build a custom MCMC sampling scheme. Details of the computational scheme are provided in Section 2.2.3.

#### 2.2.3 Computational Inference

MCMC sampling from the full posterior distributions provided in Eq. (24) for the HMM and Eq. (29) for the HMJP rely on Gibbs sampling (7, 30, 31, 35, 37, 44, 77, 78). To form a large number of samples from these posteriors, we iterate the following

1. Update 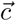 for the HMM or 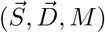 for the HMJP;
2. Update transition probabilities, that is 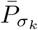 for the HMM or 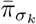 and 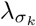 for the HMJP;
3. Update the initial probability vector 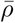 for both the HMM and the HMJP;
4. Update photon emission rates 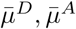 for both the HMM and the HMJP.

These samples allow us to reconstruct the posterior distribution 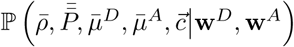 for the HMM and 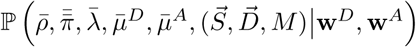 for the HMJP. The estimation of switching rates 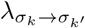 with the HMJP is carried out by Eq. (2) and by Eq. (13) for the HMM.

In Supporting Material (A), we present the equation summaries for both HMJP and HMM formulations. A working code of the the implementation of our HMM and HMJP frameworks is made available through the authors’ website.

### 2.3 Experimental Methods

Here, we introduce experimental methods. We start from sample preparation, next we move to presenting experimental procedure.

#### 2.3.1 Acquisition of Experimental Data

##### Sample Preparation

The Holliday Junction strands used in this work, and whose results are shown in Figs. 3 to 7, were purchased from JBioS (Wako, Japan), of which sequences are given below

- R-strand: 5’-CGA TGA GCA CCG CTC GGC TCA ACT GGC AGT CG-3’
- H-strand: 5’-CAT CT**T** AGT AGC AGC GCG AGC GGT GCT CAT CG-3’
- X-strand: 5’-biotin-TCTTT CGA CTG CCA GTT GAG CGC TTG CTA GGA GGA GC-3’
- B-strand: 5’-GCT CCT CC**T** AGC AAG CCG CTG CTA CTA AGA TG-3’.

We note that **T** (H-strand) and **T** (B-strand) indicate thymine residues labeled with a FRET donor (ATTO-532) and an acceptor (ATTO-647N) fluorophores, respectively, at position 6 from the 5’ end. The R, X, and B-strands (1 *μ*M, 30 *μ*L) and H-strand (1 *μ*M, 20 *μ*L) were mixed in TN buffer (10 mM Tris with 50 mM NaCl, pH 8.0). The mixture was annealed at 94°C for 4 min, and then gradually cooled down (2-3°C/min) to room temperature. We used a sample chamber (Grace Bio-Labs SecureSeal, GBL621502) and a coverslip that is coated by Biotin-PEG-SVA (Biotin-poly(ethylene glycol)-succinimidyl valerate) (79). Streptavidin (0.1 mg/mL in TN buffer, 100 *μ*L) was incubated for 20 min, which was followed by washing with TN buffer. The HJ solution (10 nM, 100 *μ*L) was injected for 3 s. The chamber was rinsed three times by measuring buffer (TN buffer with 10 mM MgCl2 and 2 mM Trolox).

##### Experiments

Broadband light, generated by super continuum laser (Fianium SC-400-4, *f* = 40 MHz), was filtered by a bandpass filter (Semrock FF01-525/30-25) and focused on the upside coverslip surface using an objective lens (Nikon Plan Apo IR 60×, N.A. = 1.27). The excitation power was set to be 20 *μ*W at the entrance port of the microscope. The fluorescence signals were collected by the same objective lens and guided to detectors through a multimode fiber (Thorlabs M50L02S-A). Fluorescence signals of donor and acceptor were divided by a dichroic mirror (Chroma Technology ZT633rdc) and filtered by bandpass filters (Semrock FF01-585/40-25 for donor and FF02-685/40-25 for acceptor), and then detected by hybrid detectors (Becker&Hickl HPM-100-40-C). For each photon signal detected, the routing information was appended by a router (Becker&Hickl HRT-41). The arrival time of the photon was measured by a Time-Correlated Single Photon Counting (TCSPC) module (Becker&Hickl SPC-130-EM) with time tagging mode.

## 3 Results

In order to show how HMJPs work and to highlight the HMJPs’ advantages over HMMs, we initially benchmark our method using simulated data that mimics smFRET experiments. Simulated data are ideal for this purpose because they have a “ground truth”. Generation of such data relies on the Gillespie algorithm (52). Next, we compare the strength of our HMJP method to HMMs on experimental data.

We focus on the following simulated dataset: a molecule that exhibits fast kinetics as compared to the data acquisition rate, see Fig. 1. Analysis of datasets on slow kinetics is relegated to Supporting Material (A). The results corresponding to the simulated dataset with fast kinetics for both HMJP and HMM are shown in Fig. 2. Subsequently, we show the performance of our method on experimental dataset Fig. 3. We present the results for the experimental data set in Figs. 4 to 7. Further analysis on different experimental datasets is also provided in Supporting Material (A).

**Fig. 2.**
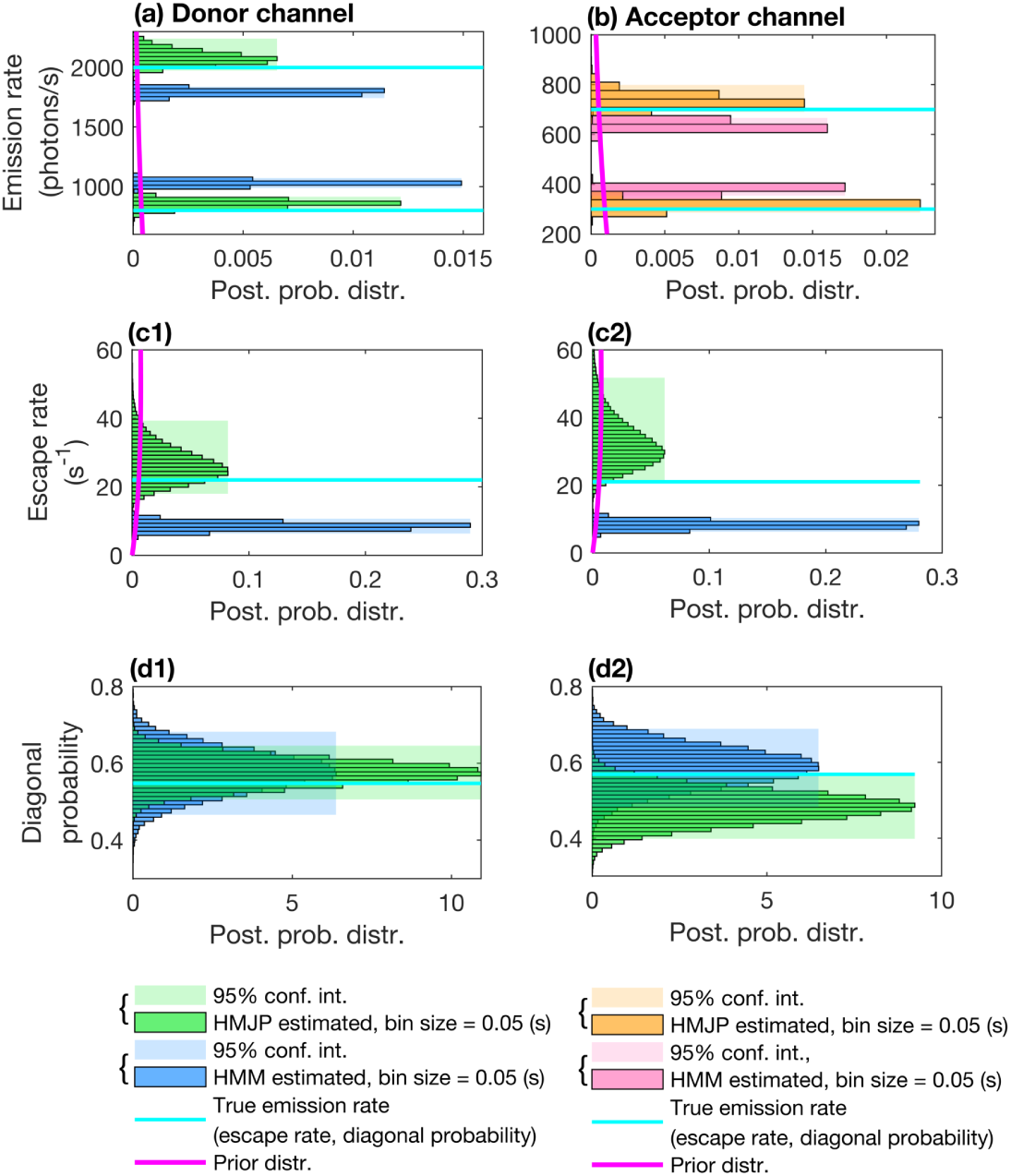
HMJP and HMM photon emission rate, escape rate and transition probability estimates for fast switching kinetics in simulated measurements. Here, we provide posterior photon emission rate, escape rate and transition probability estimates obtained with HMJP and HMM when the switching rate is faster than the data acquisition rate 1/Δt = 20 (1/s). We expect HMMs to perform poorly in estimating the true photon emission rates, escape rates and transition probabilities when the system switching is fast. In this figure’s panels (a)-(b), we superposed the posterior distributions over photon emission rates for HMJP (green for donor, orange for acceptor) and HMM (blue for donor, pink for acceptor) along with their 95% confidence intervals and the true photon emission rates (dashed cyan line). Next, in panels (c1)-(c2) and (d1)-(d2), we superposed the posterior distributions over escape rates and transition probabilities (green for HMJP and blue for HMM) along with their 95% confidence intervals, true escape rates and true transition probabilities (dashed cyan lines), respectively. In this figure’s panels (c1)-(c2) correspond to the posterior distribution of escape rates that are labeled with 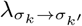 for all k, k^t^ = 1, 2 with k ≠ k^t^. Panels (d1)-(d2) correspond to the transition probabilities labeled as 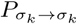 for all k = 1, 2. Here, simulated measurements are generated with the same parameters as those provided in Fig. 1 panels (b1)-(b2).

**Fig. 3.**
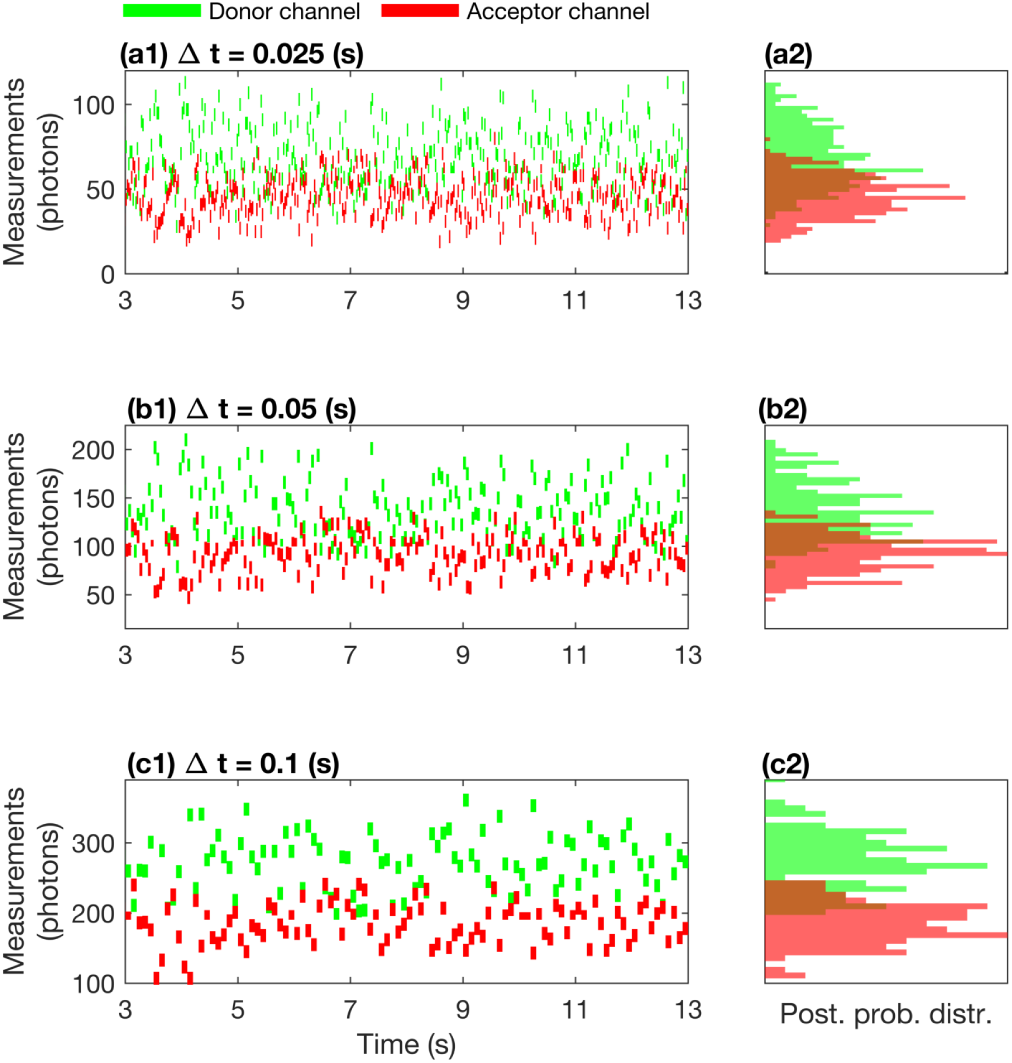
An illustration of experimental smFRET measurements for data acquisition periods. Δt = 0.025, 0.05 s and Δt = 0.1 s. In panels (a1), (b1) and (c1), we provide the measurements for the same smFRET experiment coinciding with the data acquisition periods Δt = 0.025, 0.05 s and Δt = 0.1 s. Here, we assume that the measurements are acquired by detectors with fixed exposure periods coinciding with the data acquisition periods τ = 0.025, 0.05 s and τ = 0.1 s in donor (green) and acceptor (red) channels. Panels (a2), (b2) and (c2) are especially interesting to our analysis. Here, we don’t observe well separated histograms due to fast switching kinetics of the molecule as shown in Fig. 1 panel (b2) for simulated experiments.

**Fig. 4.**
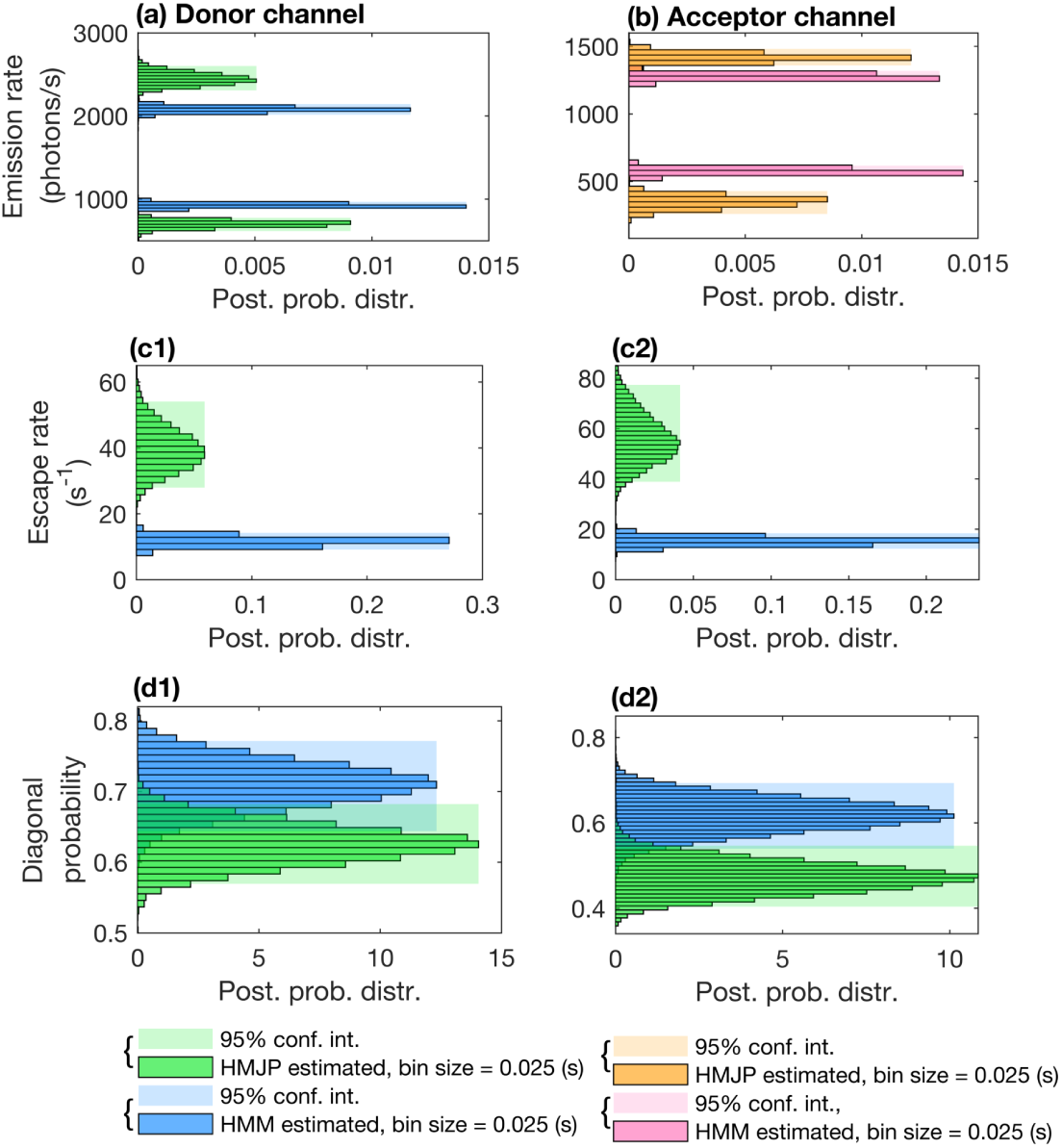
HMJP and HMM photon emission rate, escape rate and transition probability estimates for experimental smFRET measurements with. Δt = 0.025 s. Here, we provide posterior photon emission rate (panels (a)-(b)), escape rate (panels (c1)-(c2)) and transition probability (panels (d1)-(d2)) estimates obtained with HMJP and HMM for the experimental data shown in Fig. 3 panel (a1). Here, we follow a similar color convention to that of Fig. 2. However, unlike Fig. 3, ground truth information is not available for the photon emission rates, escape rates and transition probabilities as the analysis is carried out for experimental data.

Hyperparameter values used in all analyses, as well as any other choices made are presented in Supporting Material (A). For clarity, we only have access to the data demonstrated with the green and red dashes of panels (a1) and (b1) of Fig. 1 not the cyan (ground truth) trajectories. These trajectories are unknown and to be determined along with other model parameters.

### 3.1 Simulated Data Analysis

#### 3.1.1 Acquisition of Simulated Data

In the generation of our simulated data, we assumed *K* = 2 attainable states, such as on/off or folded/unfolded states for illustrative purposes only (our method trivially generalizes to more states). We assumed photon emission rates which we set at 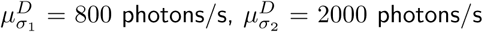 and 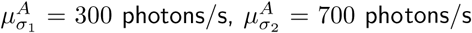 where 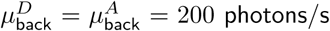. Additionally, we defined a data acquisition period of Δ*t* = 0.05 s and consider the exposure period by setting *τ* equal to 100% of Δ*t*. The onset and concluding time of the simulated data are at *t*_0_ = 0.095 s and *t*_*N*_ = 10.05 s, respectively.

We use the following structure for the switching rates 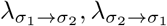 in order to specify system kinetics, with a parameter *τ*_*f*_ that sets the system kinetics time scale,

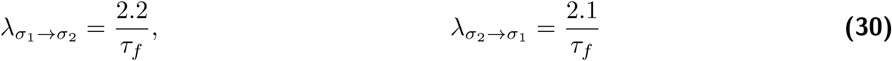

in the analysis of fast switching rates. We simulate a case with *τ*_*f*_ = 0.1 s in Fig. 1 panel (b1), which involves system kinetics that are faster than the data acquisition rate.

#### 3.1.2 Comparison of HMJPs with HMMs on Simulated Data

Here, we compare HMJPs and HMMs on the analysis of simulated measurements given in Fig. 1 panel (a1).

We expect HMJPs to perform better than HMMs as the simulated data are generated with switching rates 5% faster than the data acquisition rate (and thus many transitions occur during the data acquisition time). In this regime of switching rates, the approximation in Eq. (13) required by HMMs fails.

Given the measurements, we estimate the posterior distribution over the quantities of interest including the trajectory, 𝒯 (·), initial transition probability matrix, 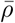, transition probabilities 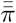 (and 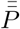 in HMM), photon emission rates, 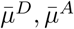 and escape rates, 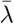. To achieve these estimates, we use HMJP and HMM samplers as introduced in Section 2.2.3 to generate pseudorandom numbers from the posterior distributions 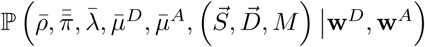 and 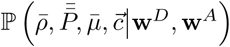, respectively.

We first show emission and escape rates estimates in Fig. 2. In Fig. 2 panels (a)-(b), (c1)-(c2), (d1)-(d2), we observe that the ground truths for photon emission rates, escape rates and transition probabilities lie within the 95% credible intervals for the corresponding HMJP estimates. By contrast, the HMM performs poorly due to the failure of the approximation in Eq. (13) resulting in unsatisfactory photon emission rate and escape rate estimates.

In particular, from Fig. 2 panels (a)-(b), we see that the HMM overestimates (by about 12% namely HMM provides estimates that are approximately 1.12 times the ground truth values for 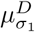 and 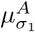) 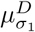 and 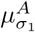 and underestimates (about 5%) 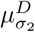 and 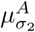. The failure of the HMM is more pronounced when looking at 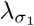 and 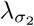. For example, in Fig. 2 panels (c1)-(c2), the HMM provides narrow distributions over escape rates by grossly underestimating both 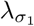 and 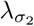 (by about 50%). On the other hand, in Fig. 2 panels (c1)-(c2), we see that HMJP posterior distribution mode coincides with the ground truth.

The HMM’s inability to capture fast kinetics is also reflected by its wide posterior distributions over transition probabilities 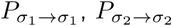 as shown in Fig. 2 panels (d1)-(d2).

In Supporting Material (A) Fig. S1, we provide the analysis on slow switching kinetics for the HMJP and HMM estimate comparisons as in Fig. 2 where both HMJP and HMM perform well.

Subsequently, in Supporting Material (A) Fig. S2, we provide the robustness of HMJP posterior estimates over photon emission rates, escape rates and transition probabilities for the data set provided in Fig. 1 panel (b1). We observe that when the kinetic rates are more than 4 times the data acquisition rate then the HMJP starts failing in providing consistent posterior estimates over over photon emission rates, escape rates and transition probabilities.

### 3.2 Experimental Data Analysis

We now move onto the analysis of experimental data sets.

#### 3.2.1 Analysis Overview

Having benchmarked our HMJP on the simulated smFRET data set, we now move on to assessing the HMJP’s performance on experimental smFRET data provided in Fig. 3, for Holliday junctions (HJ) as described in sample preparation. We deliberately start from single photon arrival data sets. Such data sets give us the flexibility to bin data as we please in order to mimic experimental data collected over broad range binning conditions as is the case in (5, 6, 21–27). In particular, we bin using 3 data acquisition periods: Δ*t* = 0.025 s (with results shown in Figs. 4 to 7), Δ*t* = 0.05 s (with results shown in Figs. 4 to 7) and Δ*t* = 0.1 s (with results shown in Figs. 4 to 7). We pick these bin sizes for a particular reason. We estimate for the smallest bins the rates ∼ 35 − 45 (1/s). Then, we consider data acquisition rates that are twice and four times bigger. We also set the exposure period, *τ* to 100% of the Δ*t* namely *τ* = 0.025, 0.05 s and *τ* = 0.1 s. As we will see, the HMJP returns values for the rates that are consistent across bin sizes as would be expected for a method that can learn transition rates slower or faster than the data acquisition rate. By contrast, the failure of HMM becomes apparent due to the inconsistency of the estimates for the switching kinetics as a function of data bin size. The HJs we use have two well characterized conformational states (3, 80). Here, our goal is to estimate the HJ emission and switching rates in addition to the trajectory of HJs simultaneously.

In Fig. 3 panels (a1), (b1), (c1), we show the data under all 3 different bin sizes. From the data, we estimate the posterior distribution over the trajectory, 𝒯 (·), initial transition probability matrix, 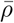, transition probabilities 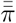 (and 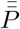 in HMM), photon emission rates, 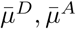 and escape rates, 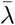. In Fig. 4, we show a direct comparison of the HMJP and HMM under the smallest bin size, where we anticipate both HMM and HMJP to be consistent with one another. As we move to larger bin sizes, the HMJP remains consistent with the result that we obtain for the smallest bin but the HMM eventually shows a lack of consistency as the bin size increases. This is apparent from Fig. 5, where we look at increasing bin size for HMM and in Fig. 6 where we look at increasing bin size for the HMJP. Finally in Fig. 7, we show the estimates over FRET efficiencies where we increase bin sizes for the case of both HMM and HMJP.

**Fig. 5.**
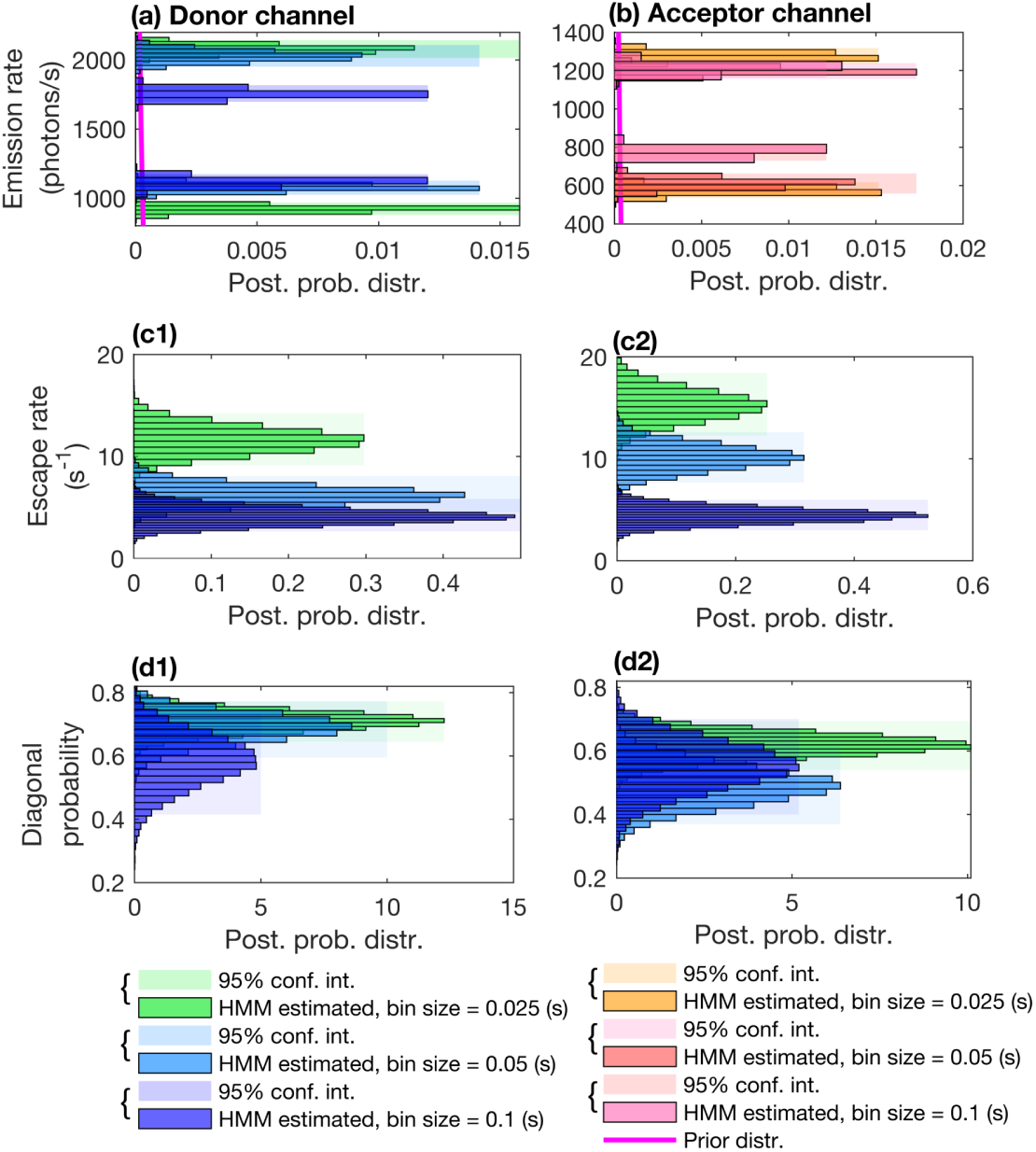
HMM photon emission rate, escape rate and transition probability estimates for experimental smFRET measurements with. Δt = 0.025, 0.05 s and Δt = 0.1 s. Here, we provide posterior photon emission rate, escape rate and diagonal transition probability estimates obtained with HMM for the measurements shown in Fig. 3 panels (a1), (b1) and (c1). We expect HMM posterior estimates associated with 3 different exposure periods not to be consistent as the switching kinetics approach the data acquisition rate. In panels (a)-(b), we superposed the posterior distributions for the measurements associated with exposure periods Δt = 0.025, 0.05 s and Δt = 0.1 s over photon emission rates for HMM along with their 95% confidence intervals. Next, in panels (c1)-(c2), we superposed the posterior distributions over escape rates along with their 95% confidence intervals. Finally, in panels (d1)-(d2), we superposed the posterior distributions over transition probabilities and their 95% confidence intervals. Posterior distributions over photon emission rate, escape rate and diagonal transition probability estimates for the integration periods Δt = 0.025, 0.05 s and Δt = 0.1 s are demonstrated in green, blue and purple, respectively.

**Fig. 6.**
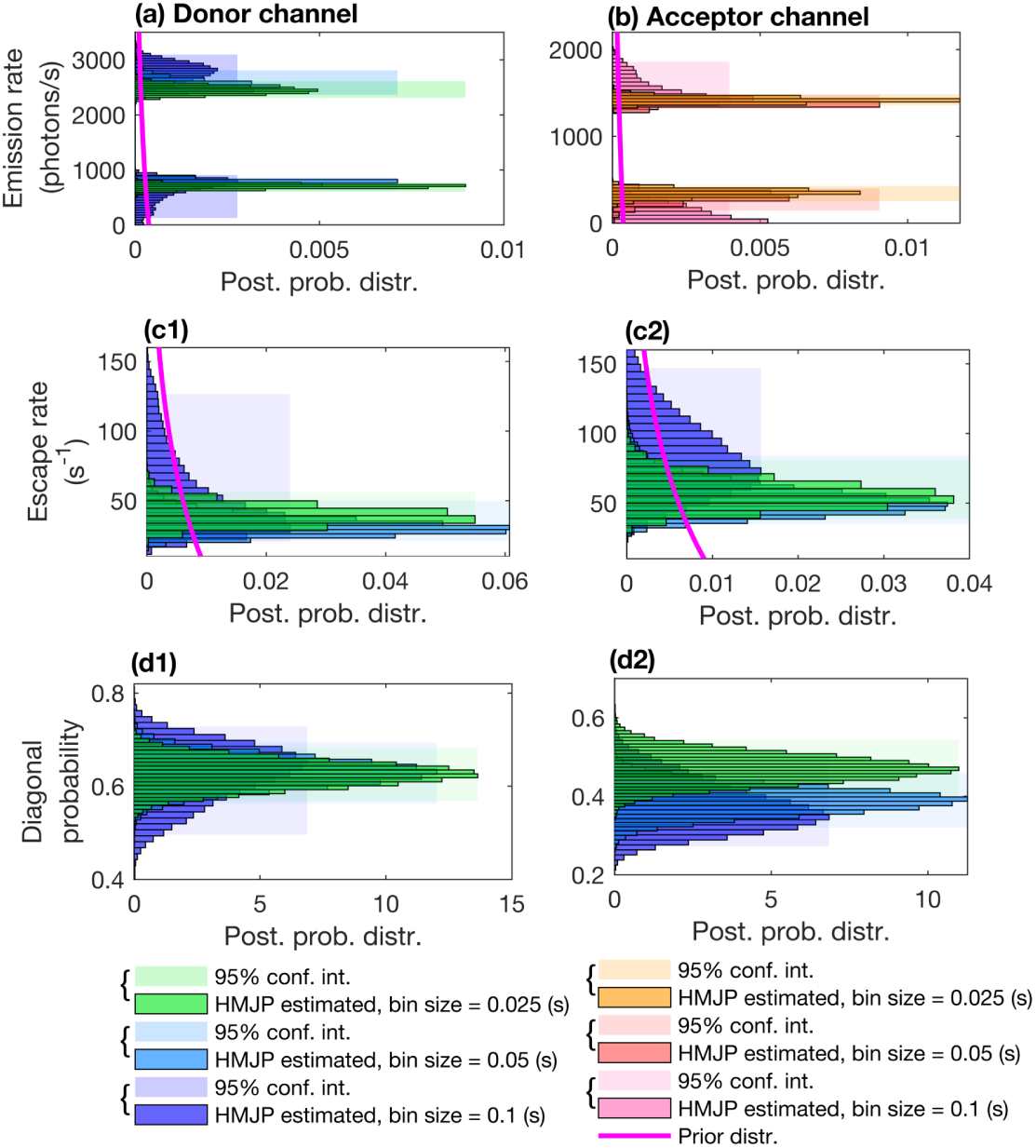
HMJP photon emission rate, escape rate and transition probability estimates for experimental smFRET measurements with. Δt = 0.025, 0.05 s and Δt = 0.1 s. Here, we provide posterior photon emission rate, escape rate and diagonal transition probability estimates obtained with HMJP for the measurements shown in Fig. 3 panels (a1), (b1) and (c1). We expect HMJP posterior estimates associated with 3 different exposure periods to be consistent across examples even if the switching kinetics exceed data acquisition rate. In panels (a)-(b), we superposed the posterior distributions for the measurements associated with exposure periods Δt = 0.025, 0.05 s and Δt = 0.1 s over photon emission rates for the HMJP along with their 95% confidence intervals. Next, in panels (c1)-(c2), we superposed the posterior distributions over escape rates along with their 95% confidence intervals. Finally, in panels (d1)-(d2), we superposed the posterior distributions over transition probabilities and their 95% confidence intervals. Here, we follow a similar color convention to that of Fig. 5.

**Fig. 7.**
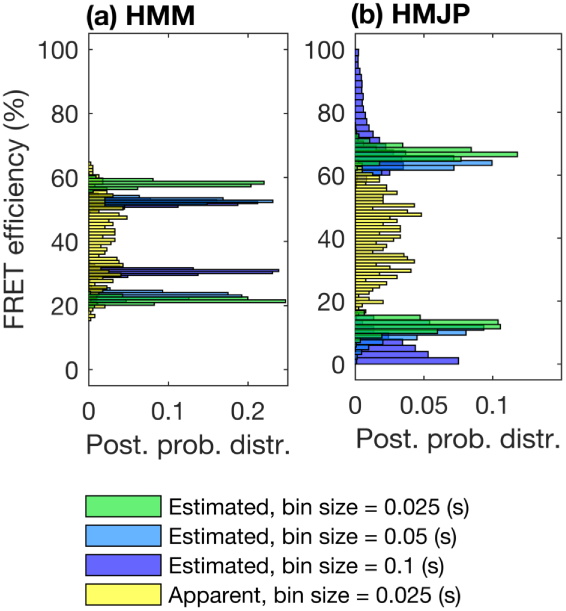
HMM and HMJP FRET efficiency estimates for experimental smFRET measurements with. Δt = 0.025, 0.05 s and Δt = 0.1 s. Here, we provide posterior FRET efficiency estimates obtained with HMM and HMJP for the measurements shown in Fig. 3 panels (a1), (b1) and (c1). We expect inconsistent posterior FRET efficiency estimates for HMMs associated with 3 different exposure periods; see panel (a). On the other hand, we expect HMJPs to provide consistent posterior estimates over FRET efficiencies for 3 different exposure periods; see panel (b). Here, in panels (a)-(b), we superposed the posterior distributions over FRET efficiencies along with the apparent FRET efficiency (yellow). In this figure, we follow a similar color convention to that of Fig. 5.

#### 3.2.2 Comparison of HMJPs with HMMs on Experimental Data

We begin with an HMJP and HMM comparison for the smallest bin size, Δ*t* = 0.025 s; see Fig. 3 panel (a1), on the photon emission rates, escape rates and transition probability estimates provided in Fig. 4. We point out that the simulated dataset provided in Fig. 1 mimic the dataset provided in Fig. 3. Even though, for experimental data Fig. 3 panels (a1), “ground truth” photon emission rates, escape rates and transition probabilities are not known, we suspect that similar observations provided in Fig. 2 hold for Fig. 4.

In Fig. 4 panels (a)-(b), we observe that HMM overestimates 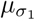 and underestimates 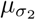 compared to HMJP estimates for 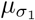 and 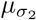. In addition, as can be seen in Fig. 4 panels (c1)-(c2), HMM provides narrow distributions over escape rates for both 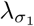 and 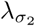. This contradicts with the escape rate posterior of the HMJP; see panels (c1)-(c2) of Fig. 4. We speculate that as HMM is unable to resolve fast switching kinetics, missed kinetic transitions are reflected in terms of underestimation/overestimation of photon emission rates or escape rates; see Fig. 4 panels (a)-(b) and (c1)-(c2). In Fig. 4 panels (d1)-(d2), we observe that HMM and HMJP estimates for 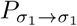 are overlapping although they have distinct posterior distributions for 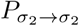.

Next, we show in Fig. 5 that the HMMs’ photon emission rates, escape rates and transition probability estimates grow inconsistent for larger bin sizes, Δ*t* = 0.05, 0.1 s (see Fig. 3 panels (b)-(c)). As a result of binning, we have multiple different apparent photon emission rates that appear. This could be overinterpreted as noise by the HMM; see panels (a)-(b). Indeed, HMMs interpret the various photon emission rates as an increased noise variance. As a related artifact, the HMM misses multiple transitions and therefore grossly underestimates escape rates.

As we see in panels (c1)-(c2), HMMs start underestimating escape rates as bin sizes become larger from Δ*t* = 0.025 s to Δ*t* = 0.05, 0.1 s. We emphasize that these escape estimates are approximate escape rate estimates obtained from the transition probability estimates for the HMM (see Eq. (13)) because a HMM does not directly report on kinetic rates. We observe in Fig. 5 panels (d1)-(d2) that the uncertainty around the HMM transition probability estimates increase and are furthermore not consistent as bin sizes increase. Subsequently, we repeat the exact same analysis as we did for the HMM for the HMJP. In particular there is an exact correspondence between Fig. 5 panels (a)-(b), (c1)-(c2), (d1)-(d2) and Fig. 6 panels (a)-(b), (c1)-(c2), (d1)-(d2). Details are provided in the captions however the important message is that HMJP escape rate estimates (see Fig. 6 panels (c1)-(c2)) has remained consistent across increased bin sizes.

Finally, in Fig. 7 panels (a), (b), we provide the posterior distributions over FRET efficiencies for the HMM and the HMJP. Here, we observe that HMMs provide inconsistent posterior FRET efficiency estimates for larger bin sizes; see panel (a). On the other hand HMJPs provide consistent FRET efficiency estimates; see panel (b).

In Supporting Material (A), we provide similar analysis for two other *experimental* smFRET data sets; see Fig. S3, Fig. S4. These analyzed data sets provided in Fig. S3, Fig. S4 were acquired under same experimental conditions as presented here. By analyzing these additional data sets, we justify the robustness of our method; see Figs. S4 to S10.

## 4 Discussion

Single molecule FRET data has the potential of revealing switching kinetics occurring on time scales at or even exceeding the measurement time scale (1, 3, 10, 32). This is especially helpful for smFRET data collected in a binned fashion (5, 6, 21–27). To achieve this, we have extended the HMM paradigm, previously used in the analysis of smFRET, by treating physical processes as they occur in nature (1, 3, 5–7, 10, 21–27, 32); that is, as continuous time processes using a Markov jump process framework (15, 47, 71–75, 81, 82).

The framework that we present here can treat multiple different data collection regimes. For example, it can treat different exposure periods, it can also treat different measurement models and variable background levels. Our framework reduces to existing frameworks, such as the HMM, in specific limits. For example, when the exposure period (*τ*) is as long as the data acquisition period (Δ*t*), and the only measurements available are those precisely at the data acquisition time, then our model reduces to the HMM. Similarly, our method is more general than the H^2^MM (3) a previously developed method that treats the continuously evolving molecule trajectory by discretizing this process on a finer time scale than the data acquisition period. In the infinitely small discretization of this grid, the H^2^MM recovers our HMJP (3).

Finally, our method for smFRET data analysis goes beyond existing methods such as the H^2^MMs (3) or others (1) by not only making point estimates for the molecular trajectory, switching kinetics and photon emission rates. Rather, the Bayesian paradigm (54, 76) within which we operate allows us to compute full posterior distributions over these estimated quantities.

As a final note, our framework might also be extended in multiple different ways. Trivially, we can change the detector model of Eq. (5) and Eq. (6) to incorporate different kinds of detector models such as for sCMOS cameras used in (7, 62, 65) this is equivalent to a change of Eq. (5) and Eq. (6). In addition, we can extend our framework to account for photophysical states of the fluorophores as well as accounting for spectral cross bleeding between donor and acceptor channels (7) by changing the measurement model such that the photon emission rates are affected by the photophysical states of the fluorophores. A non trivial extension of our work is to consider a single photon arrival framework for smFRET as opposed to dealing with binned data (1, 3). Additionally, it would be possible to leverage the strengths of Bayesian nonparametrics to learn the number of conformational states within the HMJP (7, 30, 31, 35, 37, 49, 77, 78, 83). Overcoming these difficulties is the focus of future work.

## Acknowledgements

SP acknowledges support from NSF CAREER grant MCB-1719537 and NIH NIGMS (R01GM134426). TT acknowledges support from JSPS KAKENHI Grant Number JP19F19340. ASU cluster AGAVE and Saguaro are the main computational resources utilized in this study.

## Author contributions

ZK analyzed data and developed analysis tools; ZK and IS developed computational tools; WH, KI, TT contributed experimental data; ZK, IS, SP conceived research; SP oversaw all aspects of the projects.

## Author declaration

There is no competing interests.

## Supporting Material Datasets

Data sets are supplied in .txt format.

## A Supporting Material

### A.1 Comparison of HMJP and HMM for Simulated Data with Slow Switching Kinetics

**Fig. A.1.**
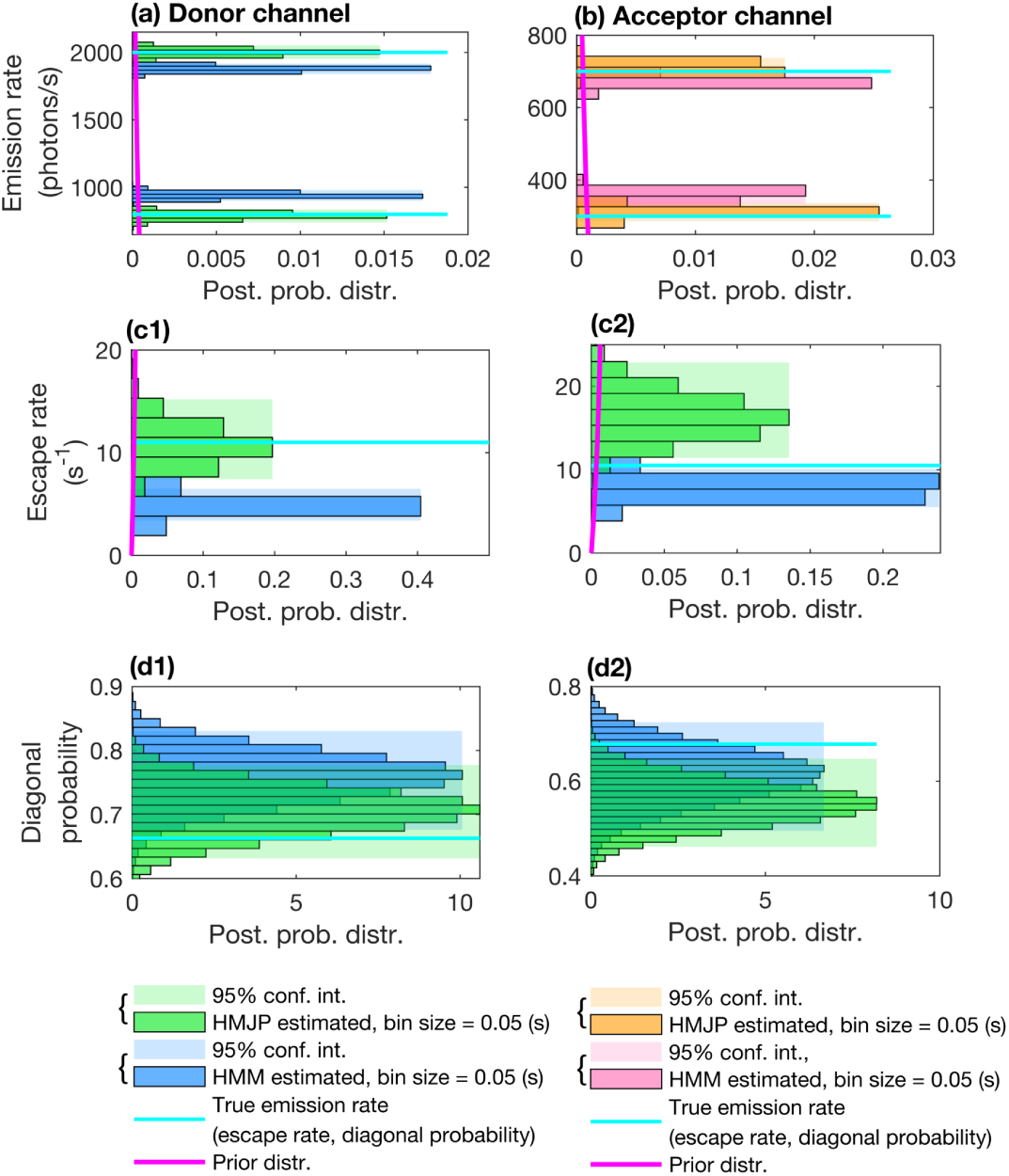
HMJP and HMM photon emission rate, escape rate and transition probability estimates for slow switching kinetics in simulated measurements. Here, we provide posteriors over the photon emission rate, escape rate and transition probability estimates obtained with HMJPs and HMMs when the switching rate is slower than the data acquisition rate 1/Δt = 20 (1/s). We expect HMMs to perform poorly in estimating the true photon emission rates, escape rates and instead perform better in estimating transition probabilities when the system switching is slow. Here, we provide the posterior distributions with the same color convention as in Fig. 2. Simulated measurements used here are generated with the same parameters as those provided in Fig. 1 panels (a1)-(a2).

### A.2 Performance of HMJP for Simulated Data with Fast Switching Kinetics over a Range of Bin Sizes

**Fig. A.2.**
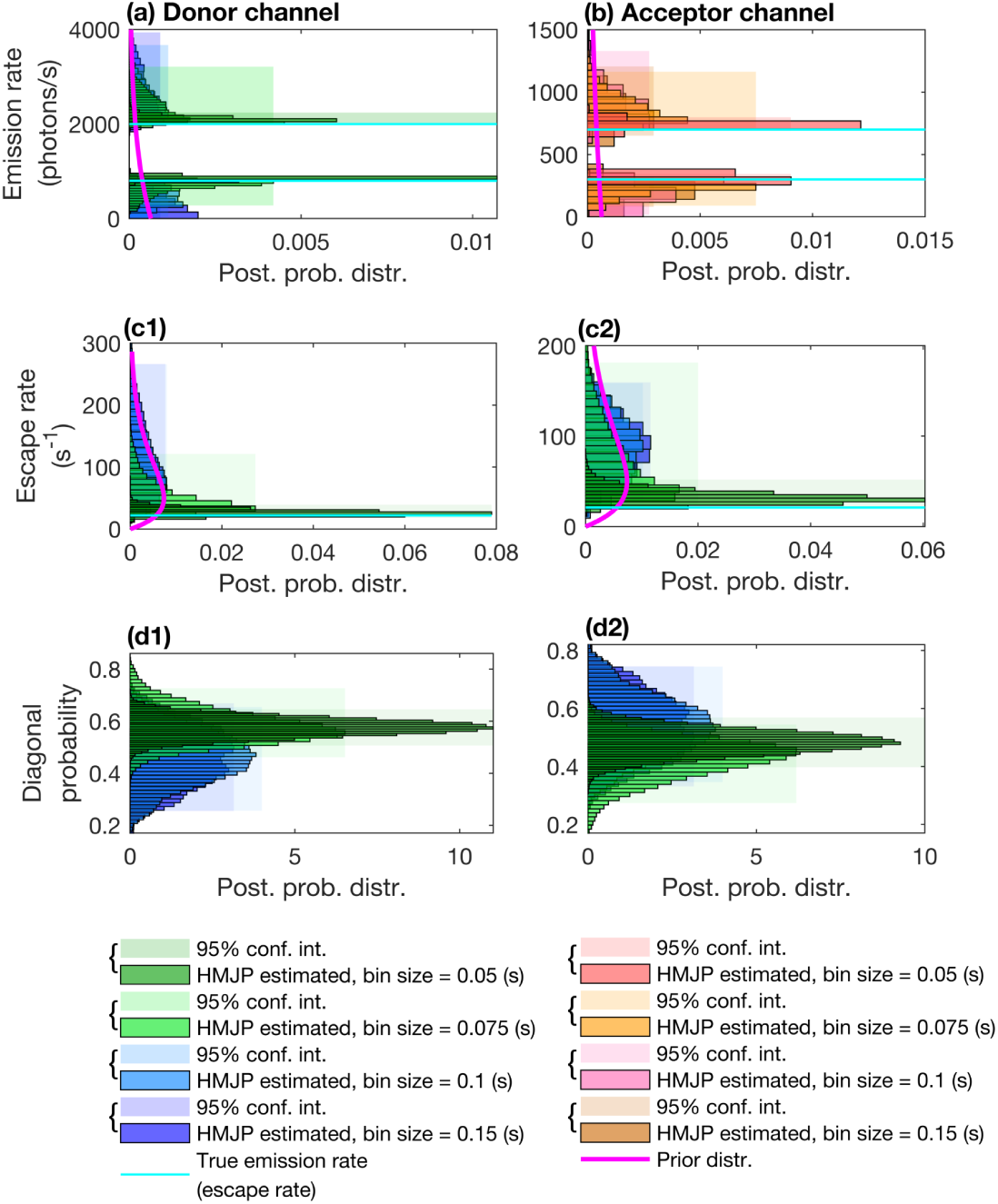
HMJP photon emission rate, escape rate and transition probability estimates for fast switching kinetics in simulated measurements. Here, we provide posterior photon emission rate, escape rate and diagonal transition probability estimates obtained with HMJP for the measurements shown in Fig. 1 panels (b1). Here, we expect HMJP posterior estimates to be consistent for the first 3 data acquisition periods (Δt = 0.05, 0.075, 0.1 s) and not consistent for Δt = 0.15 s. In this figure’s each panel, we have the same content information as in Fig. A.1. Posterior distributions over photon emission rate, escape rate and diagonal transition probability estimates for the integration periods Δt = 0.05, 0.075, 0.1 s and Δt = 0.15 s are demonstrated in dark green, green, blue and purple, respectively.

### A.3 HMJP Estimates for Additional Experimental smFRET Data

**Fig. A.3.**
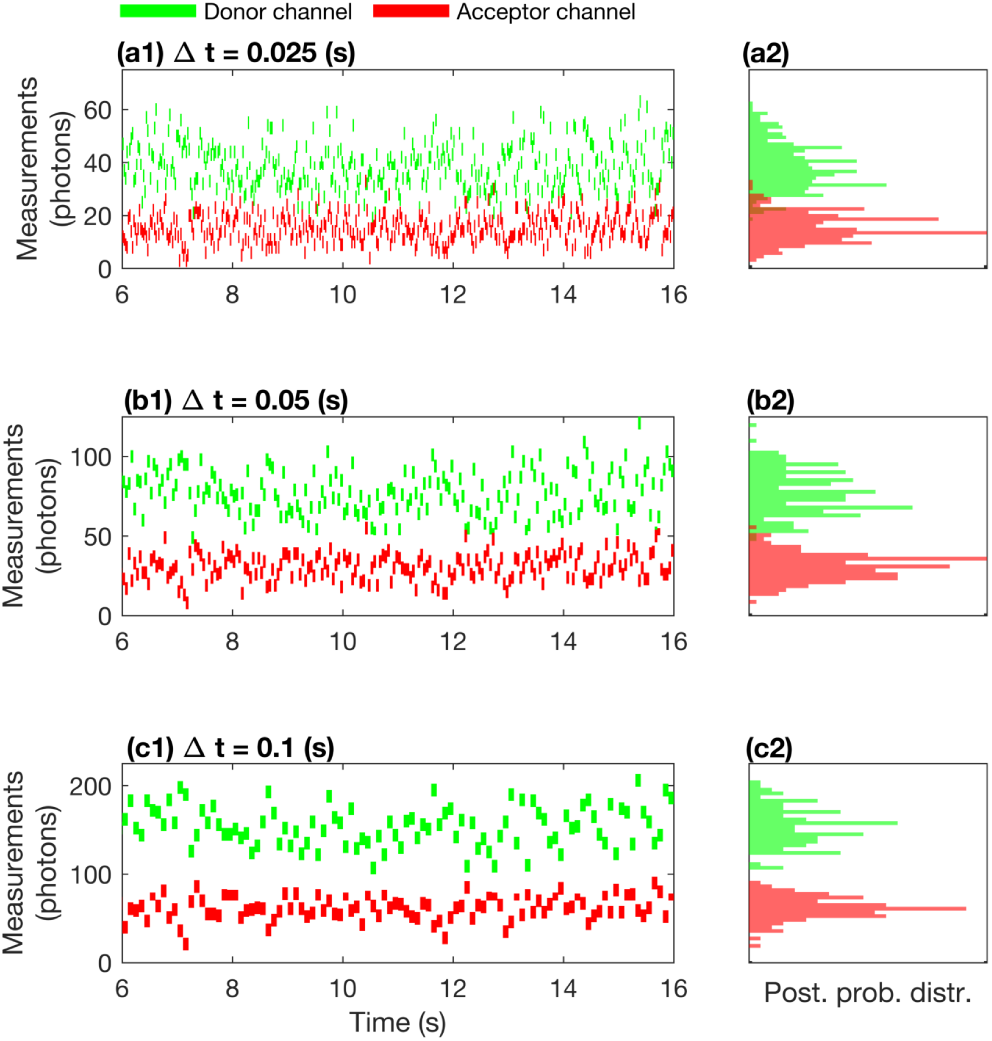
An illustration of experimental smFRET measurements for data acquisition periods. Δt = 0.025, 0.05 s and Δt = 0.1 s. In panels (a1), (b1) and (c1), we provide the measurements for the same smFRET experiment coinciding with the data acquisition periods Δt = 0.025, 0.05 s and Δt = 0.1 s. Here, we assume that the measurements are acquired by detectors with fixed exposure periods coinciding with the data acquisition periods τ = 0.025, 0.05 s and τ = 0.1 s in donor (green) and acceptor (red) channels.

**Fig. A.4.**
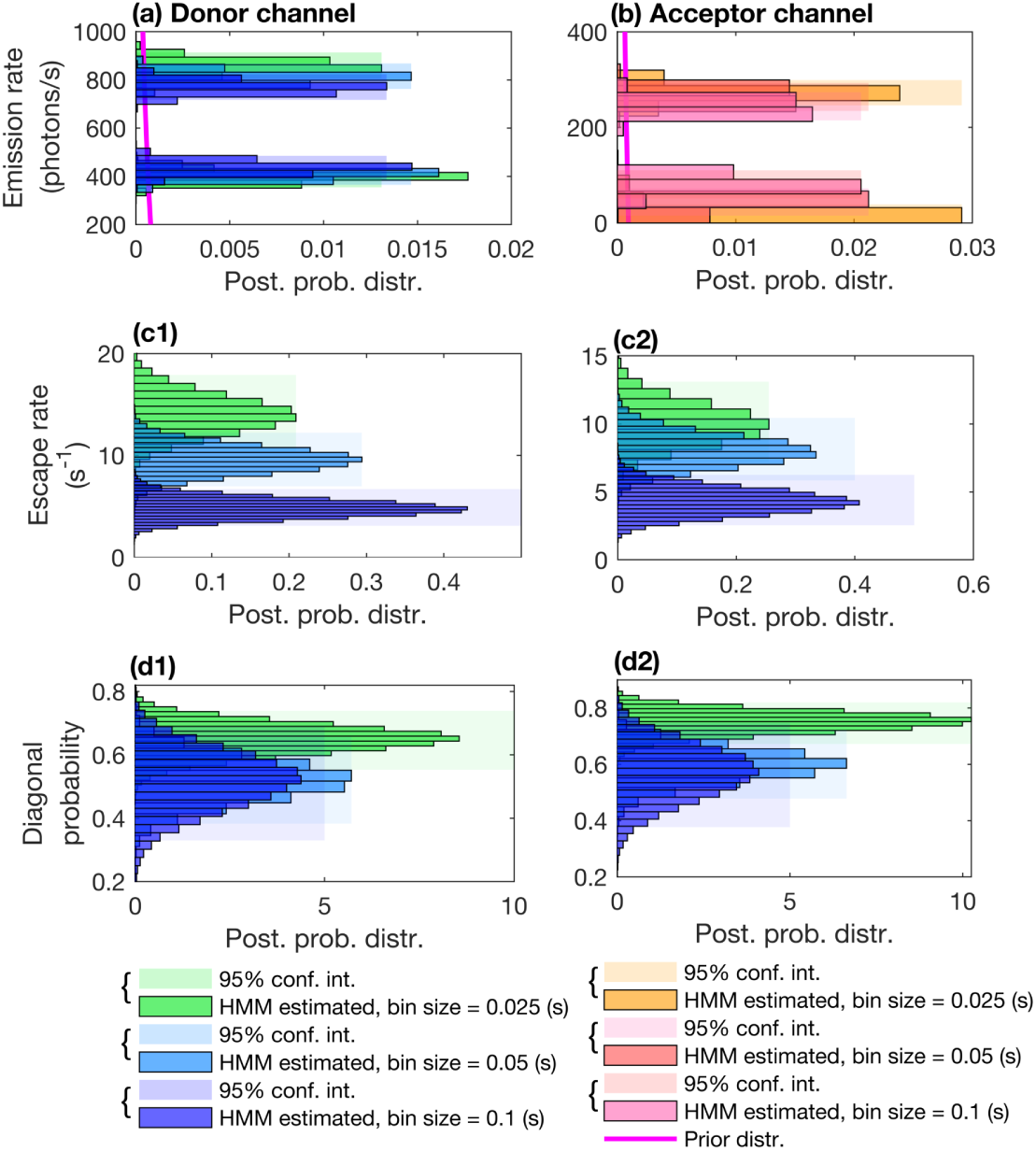
HMM photon emission rate, escape rate and transition probability estimates for experimental smFRET measurements with. Δt = 0.025, 0.05 s and Δt = 0.1 s. Here, we provide posterior photon emission rate (panels (a)- (b)), escape rate (panels (c1)-(c2)) and transition probability (panels (d1)-(d2)) estimates obtained with HMM for the experimental data shown in Fig. A.3 panels (a1),(b1) and (c1). Here, we follow a similar color convention as in Fig. 5.

**Fig. A.5.**
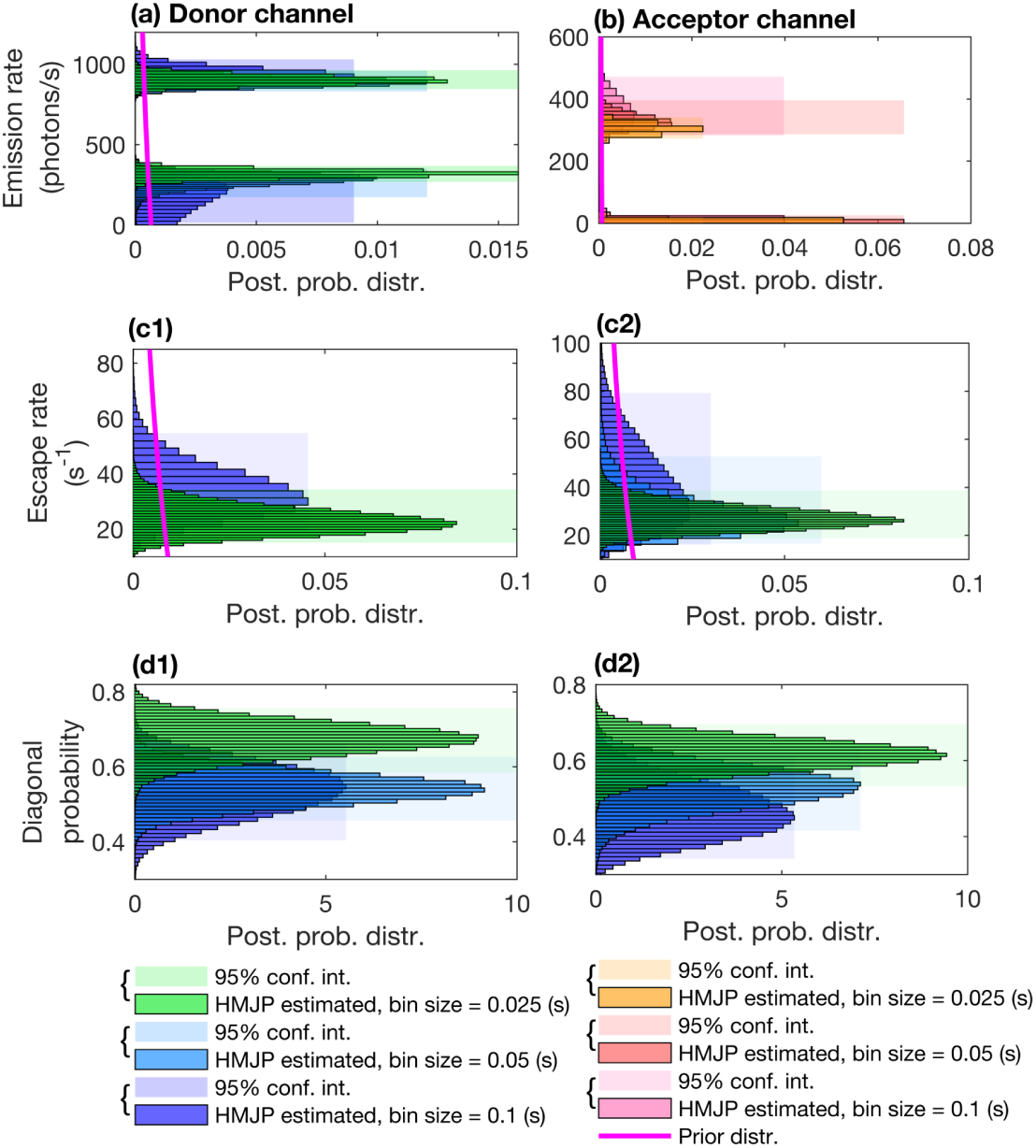
HMJP photon emission rate, escape rate and transition probability estimates for experimental smFRET measurements with. Δt = 0.025, 0.05 s and Δt = 0.1 s. Here, we provide posterior photon emission rate (panels (a)- (b)), escape rate (panels (c1)-(c2)) and transition probability (panels (d1)-(d2)) estimates obtained with HMJP for the experimental data shown in Fig. A.3 panels (a1),(b1) and (c1). Here, we follow a similar color convention as in Fig. A.4.

**Fig. A.6.**
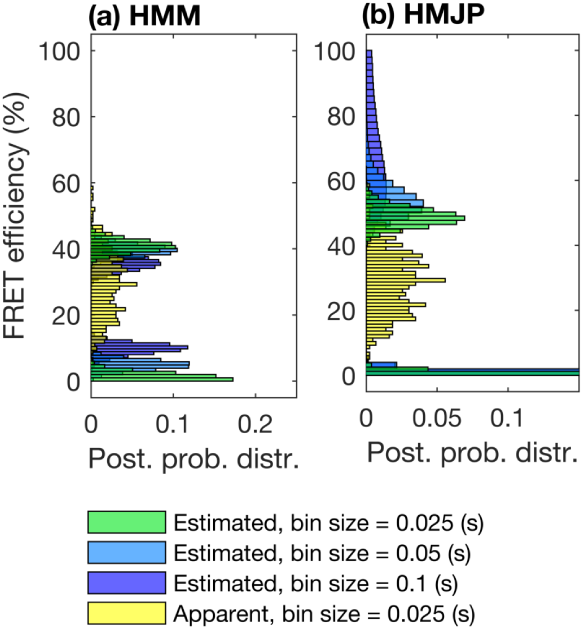
HMJP and HMM FRET efficiency estimates for experimental smFRET measurements with. Δt = 0.025, 0.05 s and Δt = 0.1 s. Here, we provide posterior photon emission rate and FRET efficiency estimates obtained with HMJP for the measurements shown in Fig. A.3 panels (a1), (b1) and (c1). We expect inconsistent posterior FRET efficiency estimates for HMM associated with 3 different exposure periods; see panel (a). On the other hand, we expect HMJP to provide consistent posterior estimates over FRET efficiencies for 3 different exposure periods; see panel (b). Here, in panels (a)-(b), we superposed the posterior distributions over FRET efficiencies along with the apparent FRET efficiency (yellow). In this figure, we follow a similar color convention to that of Fig. 5.

**Fig. A.7.**
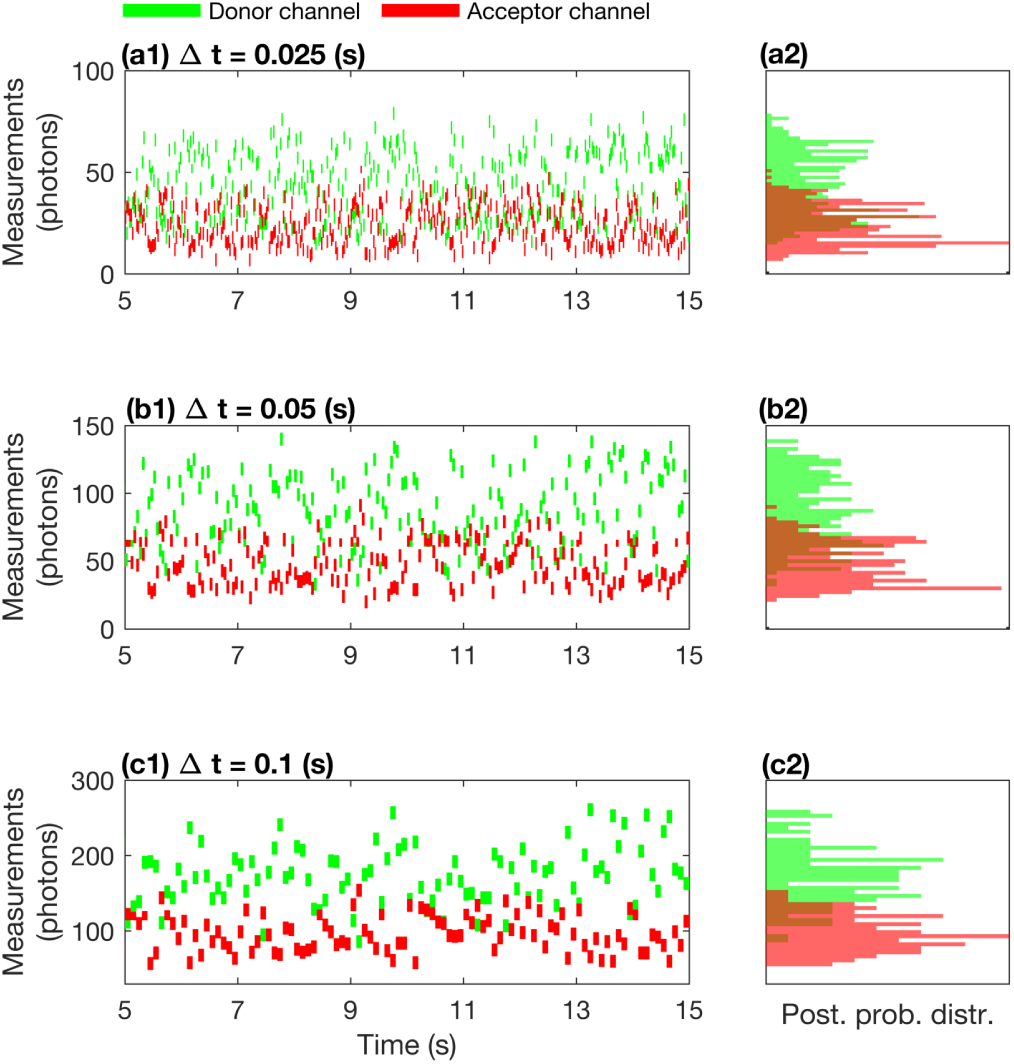
An illustration of experimental smFRET measurements for data acquisition periods. Δt = 0.025, 0.05 s and Δt = 0.1 s. In panels (a1), (b1) and (c1), we provide the measurements for the same smFRET experiment coinciding with the data acquisition periods Δt = 0.025, 0.05 s and Δt = 0.1 s. Here, we assume that the measurements are acquired by detectors with fixed exposure periods coinciding with the data acquisition periods τ = 0.025, 0.05 s and τ = 0.1 s in donor (green) and acceptor (red) channels.

**Fig. A.8.**
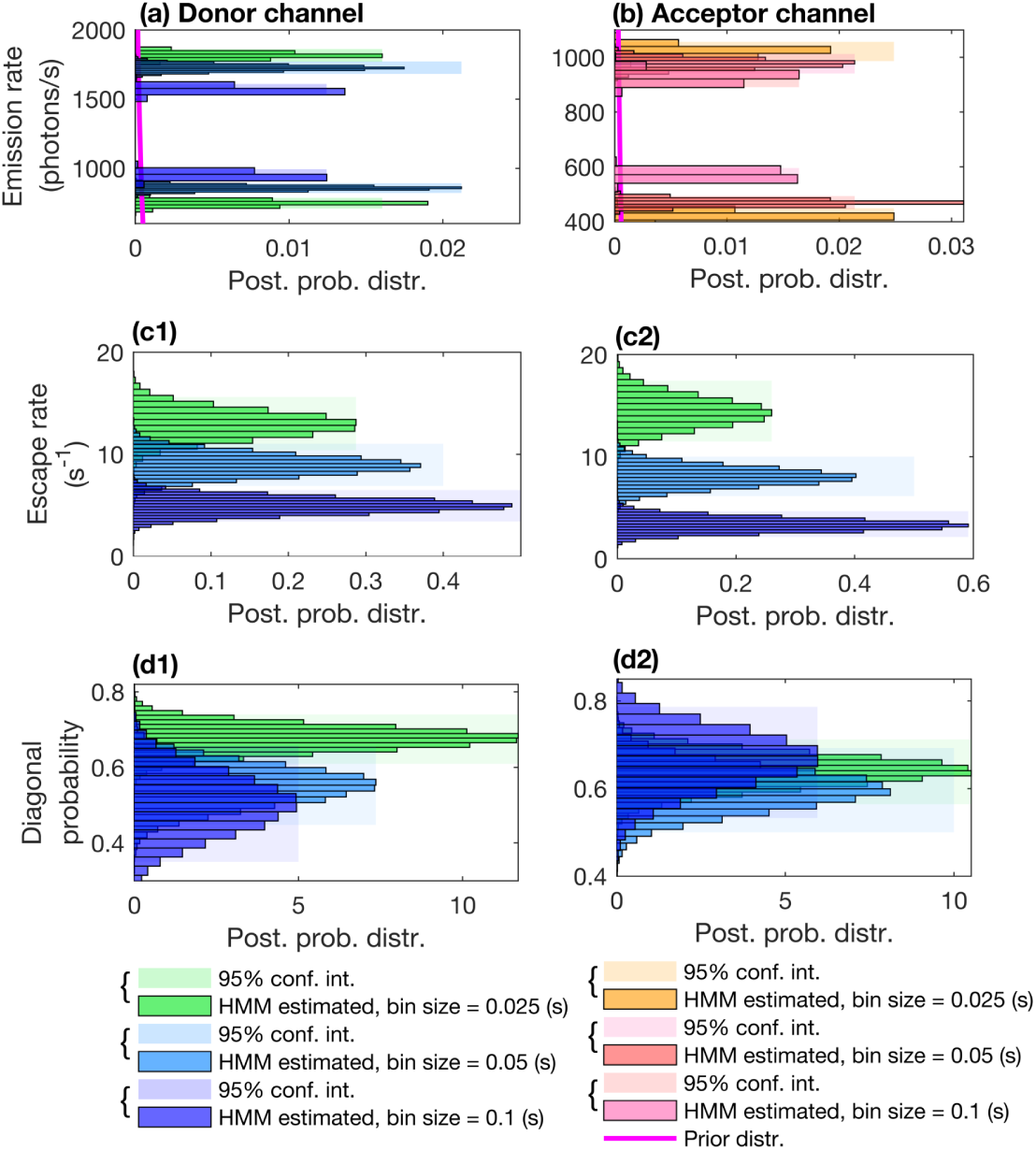
HMM photon emission rate, escape rate and transition probability estimates for experimental smFRET measurements with. Δt = 0.025, 0.05 s and Δt = 0.1 s. Here, we provide posterior photon emission rate (panels (a)- (b)), escape rate (panels (c1)-(c2)) and transition probability (panels (d1)-(d2)) estimates obtained with HMM for the experimental data shown in Fig. A.7 panels (a1),(b1) and (c1). Here, we follow a similar color convention as in Fig. 5.

**Fig. A.9.**
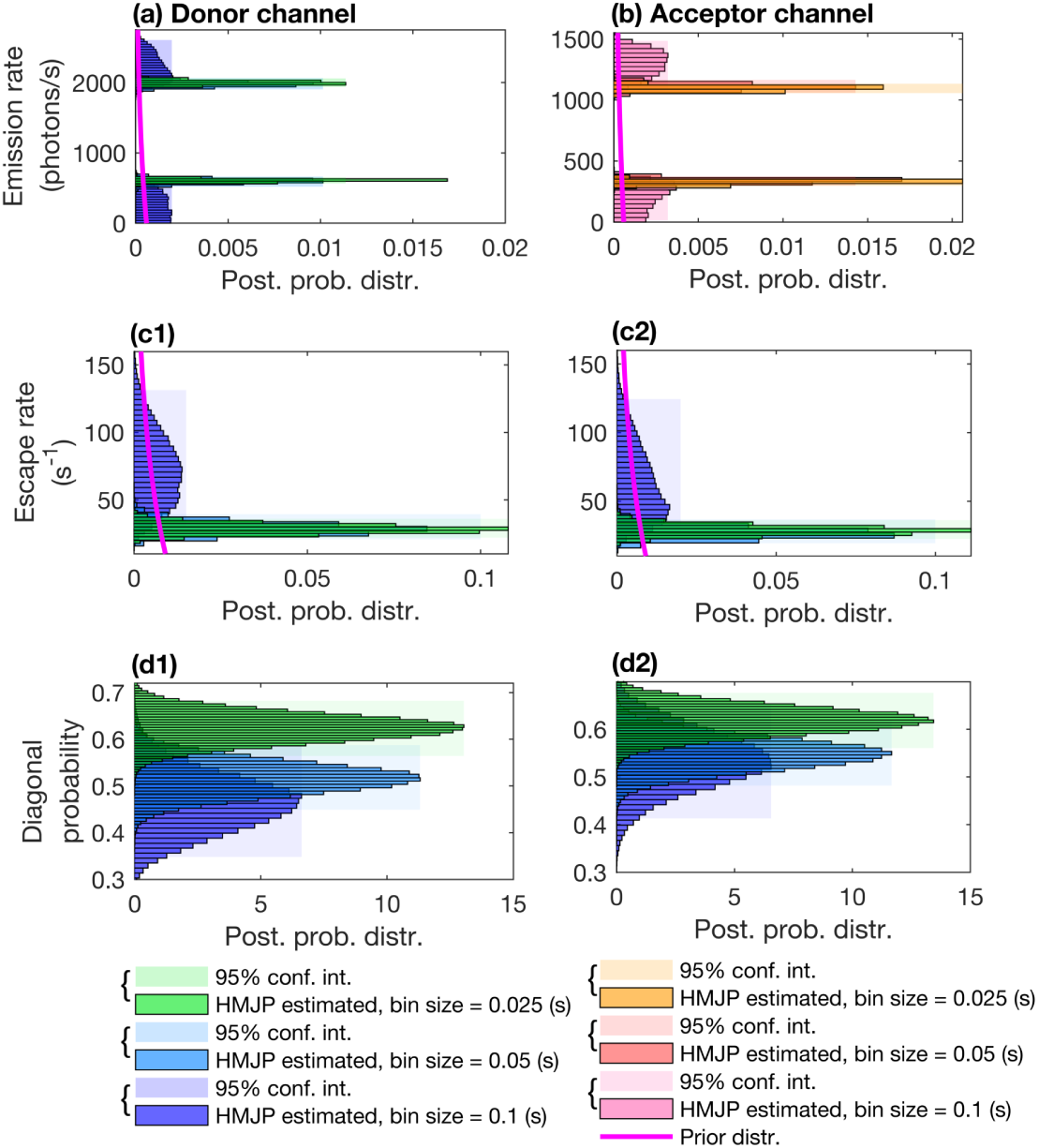
HMJP photon emission rate, escape rate and transition probability estimates for experimental smFRET measurements with. Δt = 0.025, 0.05 s and Δt = 0.1 s. Here, we provide posterior emission rate (panels (a)-(b)), escape rate (panels (c1)-(c2)) and transition probability (panels (d1)-(d2)) estimates obtained with HMJP for the experimental data shown in Fig. A.7 panels (a1),(b1) and (c1). Here, we follow a similar color convention as in Fig. A.4.

**Fig. A.10.**
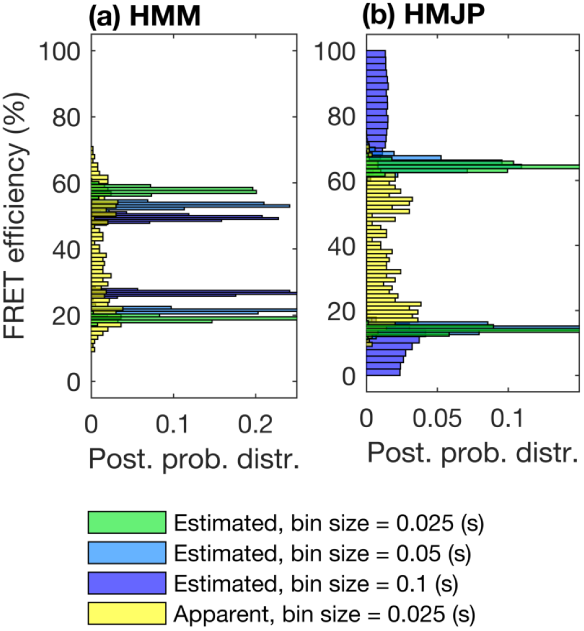
HMJP and HMM FRET efficiency estimates for experimental smFRET measurements with. Δt = 0.025, 0.05 s and Δt = 0.1 s. Here, we provide posterior photon emission rate and FRET efficiency estimates obtained with HMJP for the measurements shown in Fig. A.7 panels (a1), (b1) and (c1). We expect inconsistent posterior FRET efficiency estimates for HMM associated with 3 different exposure periods; see panel (a). On the other hand, we expect HMJP to provide consistent posterior estimates over FRET efficiencies for 3 different exposure periods; see panel (b). Here, in panels (a)-(b), we superposed the posterior distributions over FRET efficiencies along with the apparent FRET efficiency (yellow). In this figure, we follow a similar color convention to that of Fig. 5.

### A.4 Modeling Summary

#### A.4.1 Hidden Markov Jump Process Model

For *K* = 2, the full set of HMJP equations is

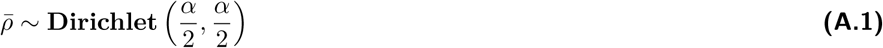

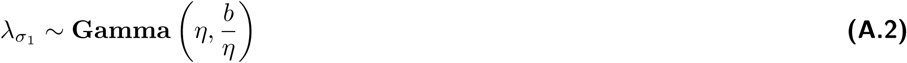

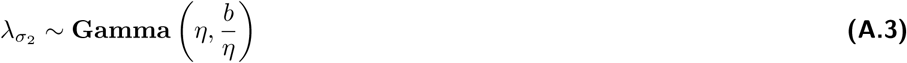

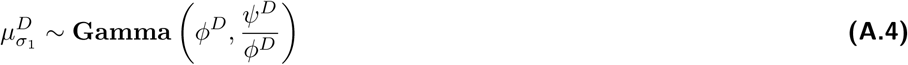

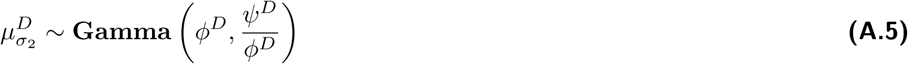

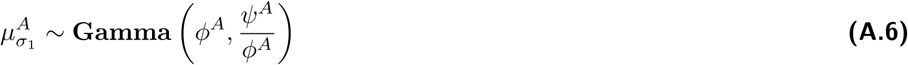

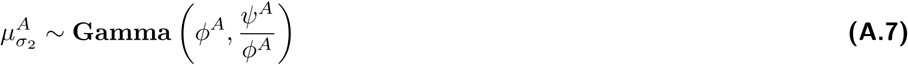

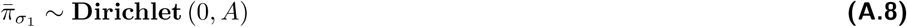

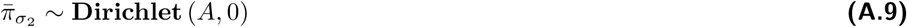

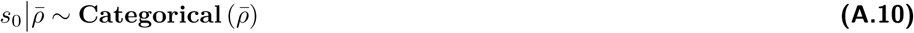

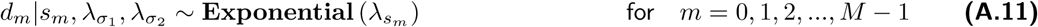

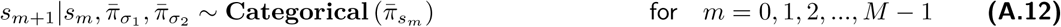

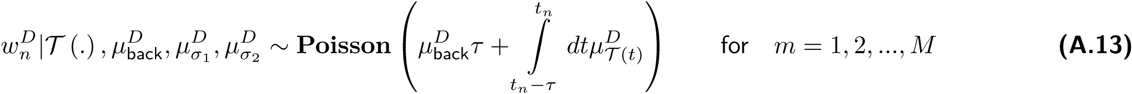

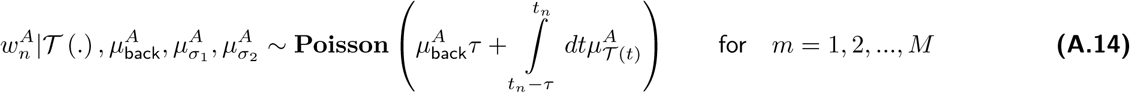

where 𝒯 (·) is formulated as follows

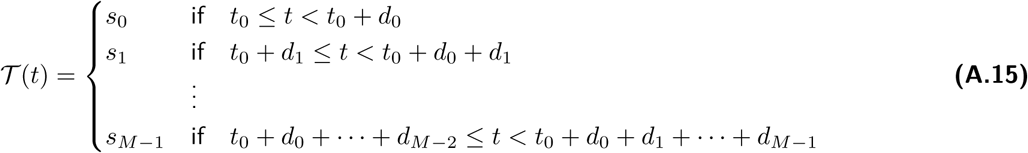

with *M* determined based on the first time

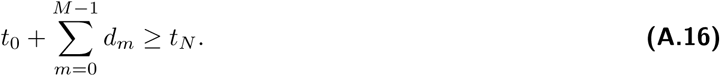

#### A.4.2 Hidden Markov Model

For *K*=2, the full set of HMM equations is

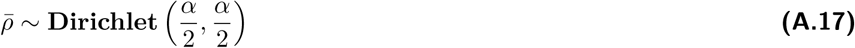

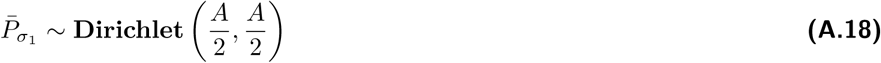

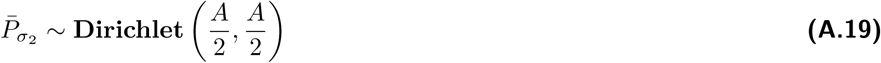

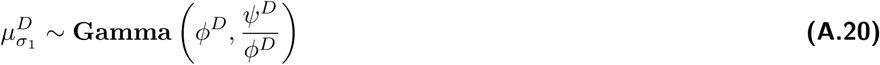

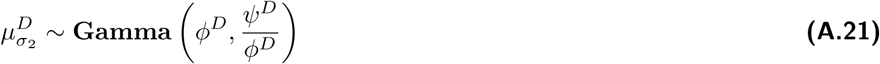

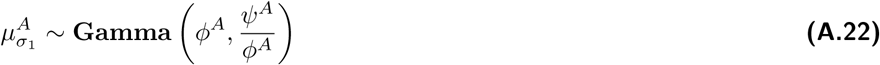

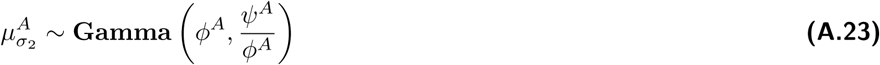

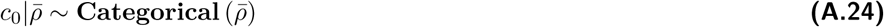

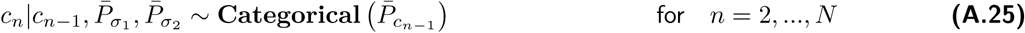

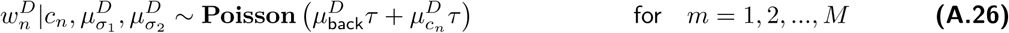

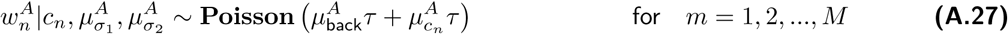

#### A.4.3 Overview of the Sampling Updates

In order to produce samples from the full posterior distributions 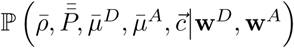 for the HMM and 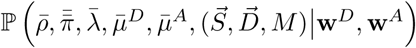 for the HMJP, we use Gibbs sampling (7, 30, 31, 35, 37, 44, 77, 78). Specifically, we repeat the following steps:

1. Update 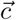 for the HMM or 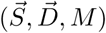 for the HMJP;
2. Update transition probabilities, that is 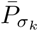 for the HMM or 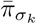 and 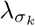 for the HMJP;
3. Update the initial probability vector 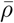 for both the HMM and the HMJP;
4. Update photon emission rates 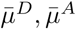 for both the HMM and the HMJP.

Here, we present the equation summaries for sampling photon emission rates.

#### Sampling photon emission rates for HMJP

In order to produce proposals for the Metropolis Hastings algorithm, we use Hamiltonian Monte Carlo (HMC). This method is very fruitful as Hamiltonian dynamics preserve volume in (*q, p*) space therefore we can use the trajectories to define complex mappings. Here the target distribution we have is the following

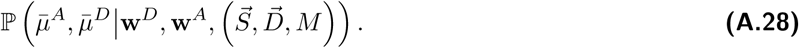

There are two main variables, these are the position that is labeled with *q* and the momentum, *p*. Here we collect the parameter of interest to be sampled in *q* such that

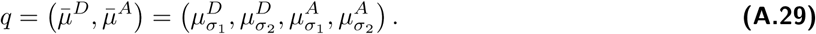

We have the following potential

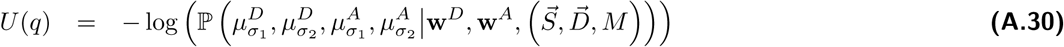

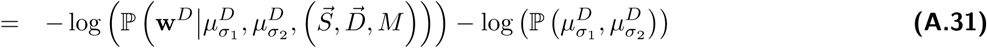

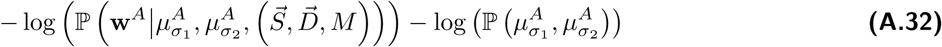

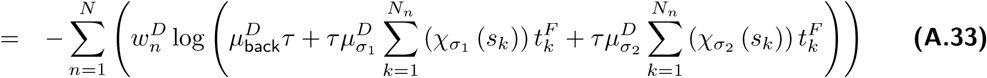

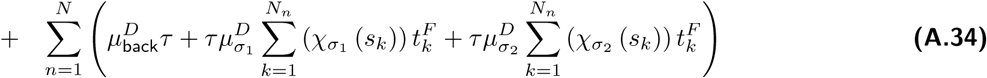

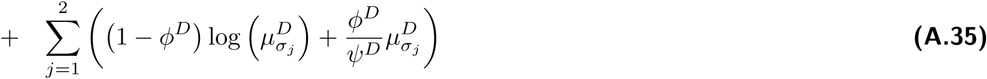

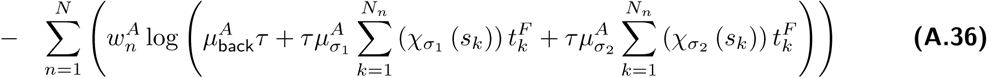

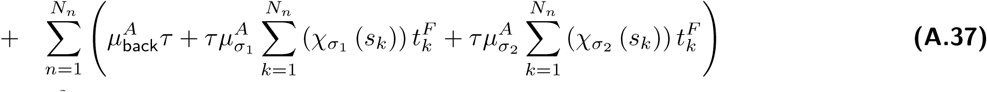

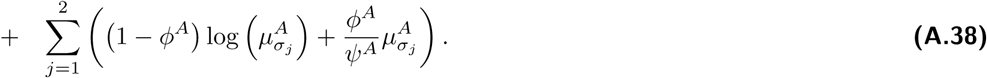

We also need the gradient of *U* (*q*) namely, 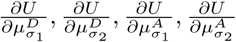.

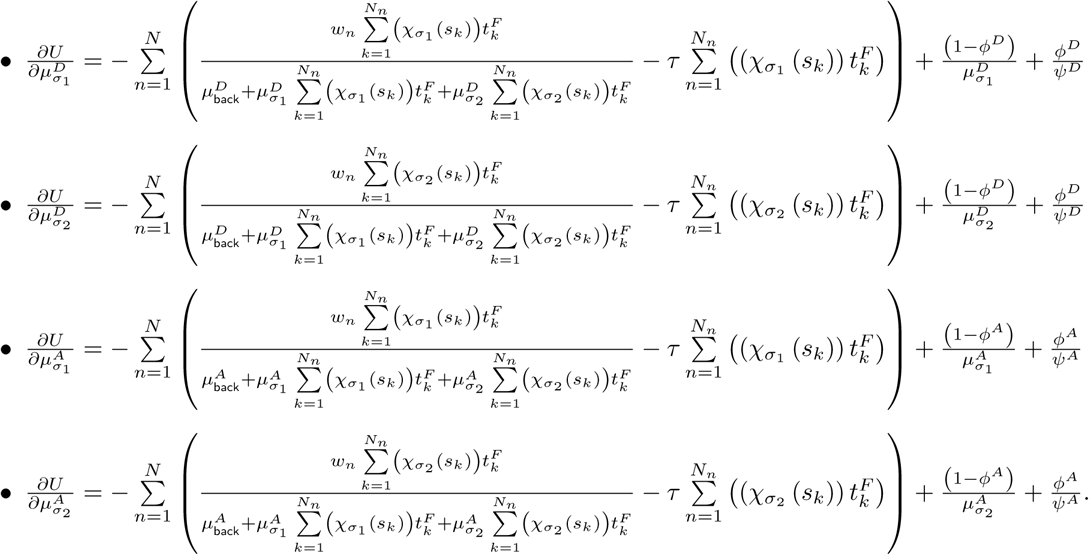

#### Sampling photon emission rates for HMM

Similarly with above, we use HMC on the target distribution

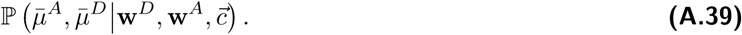

There are two main variables, these are the position that is labeled with *q* and the momentum, *p*. Here we collect the parameter of interest to be sampled in *q* such that

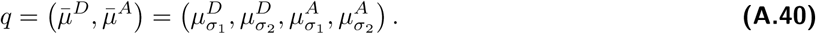

We have the following potential

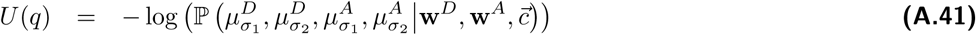

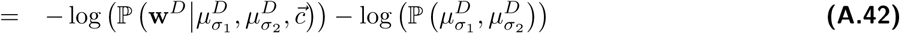

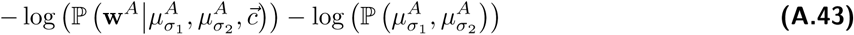

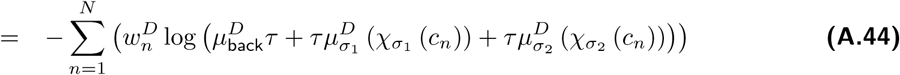

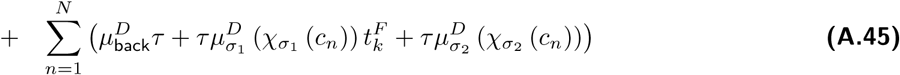

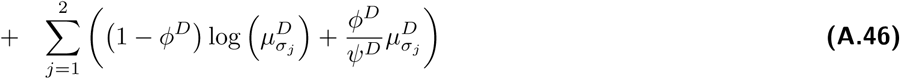

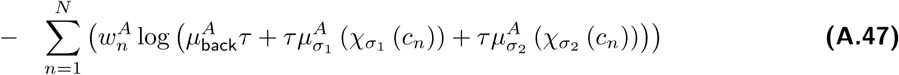

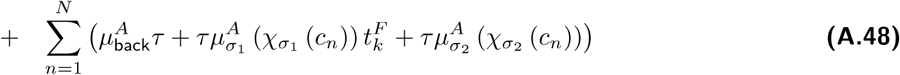

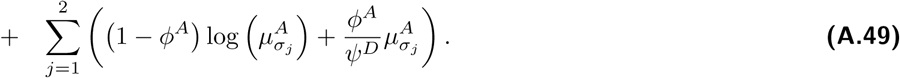

We also need the gradient of *U* (*q*) namely, 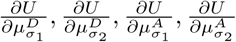.

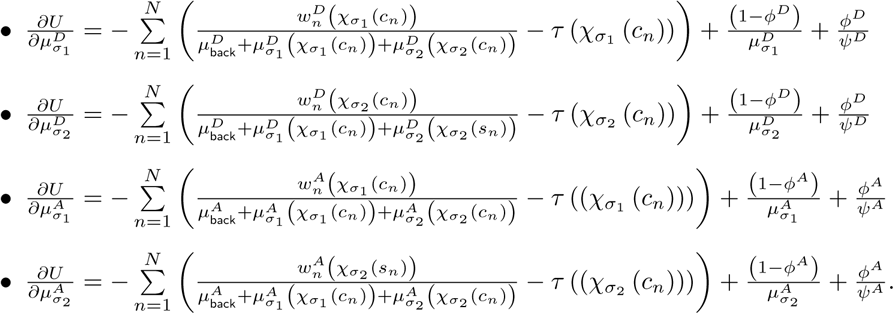

#### A.4.4 Background Photon Emission Rates

We estimate the values of 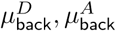 from portions of the time traces without signal, where 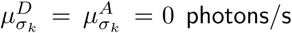. We analyze these portions, labeled with 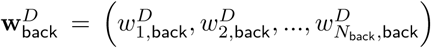 and 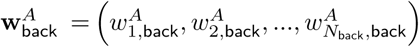, to estimate the background photon emission rates based on the formulation provided below

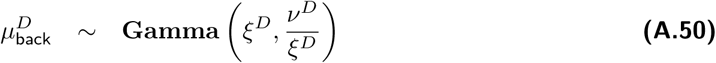

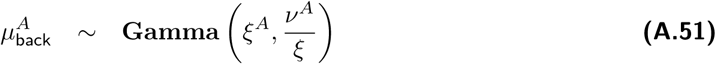

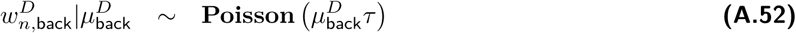

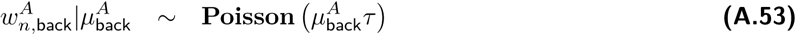

for all *n* = 1, 2, …, *N*_back_ with hyperparameters *ξ*^*D*^, *ν*^*D*^, *ξ*^*A*^, *ν*^*A*^.

Therefore, we have the following posterior distribution for the background photon emission rates for the donor channel

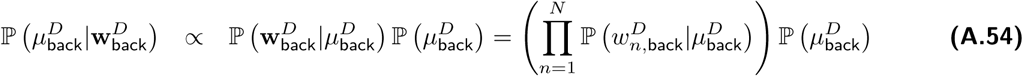

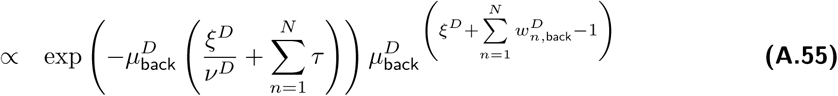

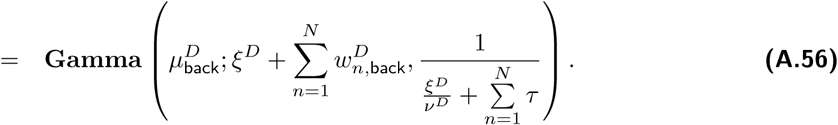

We can then estimate 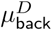 via the mean of the above posterior distribution, which is equal to 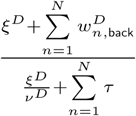. As the prior becomes non-informative (54, 76, 84); that is, at the limit *ξ*^*D*^ → 0, our estimate reduces to

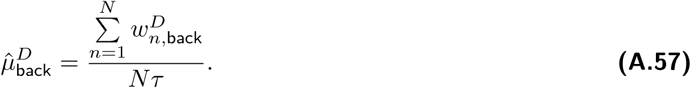

We obtain the estimator 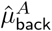 in the acceptor channel similarly.

### A.5 Notation and Analysis Options

**Table A.1.**
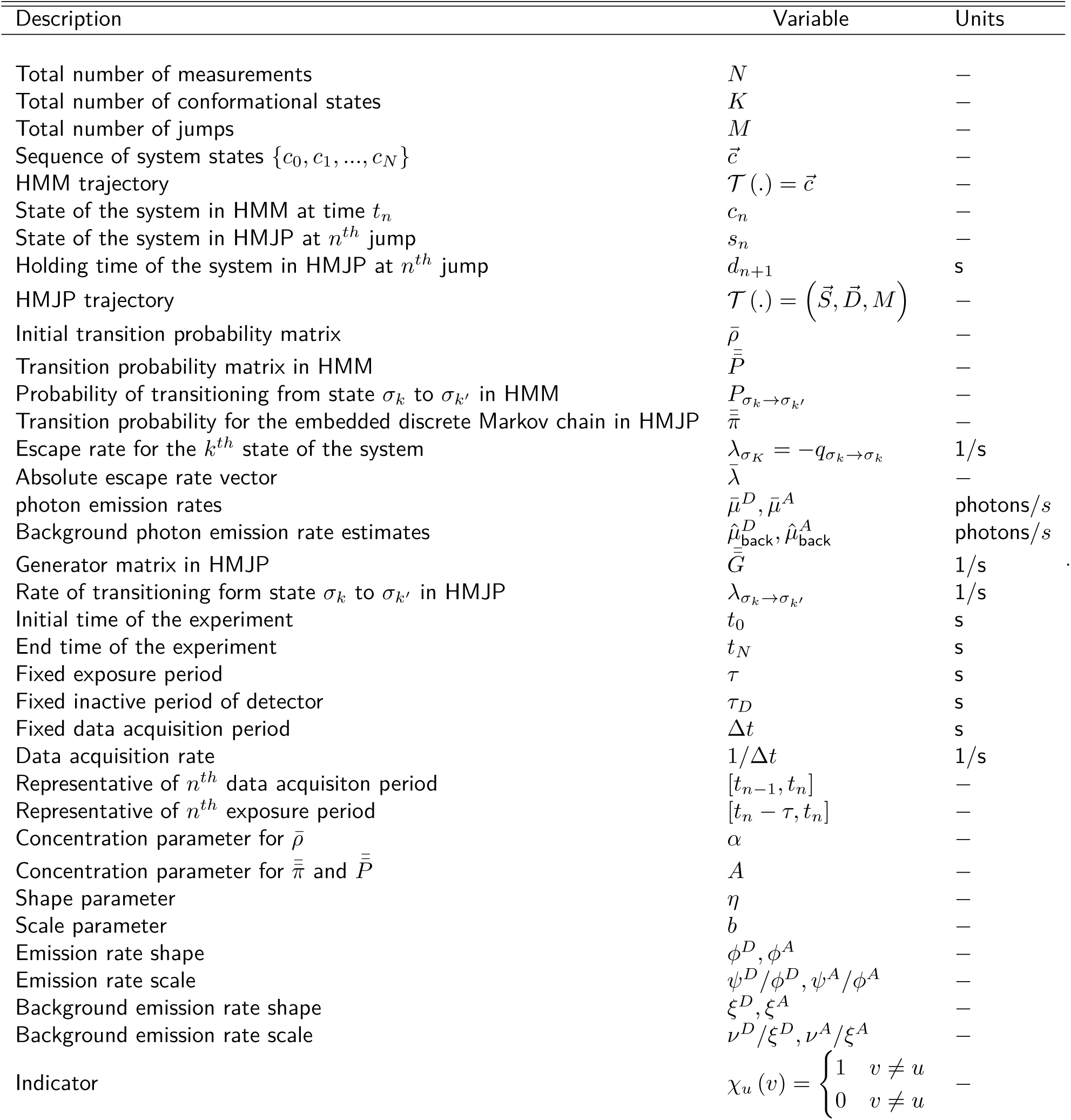
Notation Conventions.

**Table A.2.** Parameter Choices and Units.

| -<br>Units | $\psi^D, \psi^A$<br>photons/s | $\phi^D, \phi^A$<br>- | $\eta$<br>- | $b$<br>1/s | $A$<br>- | $\alpha$<br>- | $\Delta t$<br>s |
| --- | --- | --- | --- | --- | --- | --- | --- |
| Fig. 1 | total photon counts/total duration | 1 | 2 | 100 | 1 | 1 | 0.05 |
| Fig. 2 | total photon counts/total duration | 1 | 2 | 100 | 1 | 1 | 0.05 |
| Fig. 3 | total photon counts/total duration | 1 | 2 | 100 | 1 | 1 | 0.025, 0.05, 0.1 |
| Fig. 4 | total photon counts/total duration | 1 | 2 | 100 | 1 | 1 | 0.025 |
| Fig. 5 | total photon counts/total duration | 1 | 2 | 100 | 1 | 1 | 0.025, 0.05, 0.1 |
| Fig. 6 | total photon counts/total duration | 1 | 2 | 100 | 1 | 1 | 0.025, 0.05, 0.1 |
| Fig. 7 | total photon counts/total duration | 1 | 2 | 100 | 1 | 1 | 0.025, 0.05, 0.1 |
| Fig. A.1 | total photon counts/total duration | 1 | 2 | 300 | 1 | 1 | 0.05 |
| Fig. A.1 | total photon counts/total duration | 1 | 2 | 300 | 1 | 1 | 0.05 |
| Fig. A.2 | total photon counts/total duration | 1 | 2 | 300 | 1 | 1 | 0.05, 0.075, 0.1, 0.15 |
| Fig. A.3 | total photon counts/total duration | 1 | 2 | 100 | 1 | 1 | 0.025, 0.05, 0.1 |
| Fig. A.4 | total photon counts/total duration | 1 | 2 | 100 | 1 | 1 | 0.025, 0.05, 0.1 |
| Fig. A.5 | total photon counts/total duration | 1 | 2 | 100 | 1 | 1 | 0.025, 0.05, 0.1 |
| Fig. A.6 | total photon counts/total duration | 1 | 2 | 100 | 1 | 1 | 0.025, 0.05, 0.1 |
| Fig. A.7 | total photon counts/total duration | 1 | 2 | 100 | 1 | 1 | 0.025, 0.05, 0.1 |
| Fig. A.8 | total photon counts/total duration | 1 | 2 | 100 | 1 | 1 | 0.025, 0.05, 0.1 |
| Fig. A.9 | total photon counts/total duration | 1 | 2 | 100 | 1 | 1 | 0.025, 0.05, 0.1 |
| Fig. A.10 | total photon counts/total duration | 1 | 2 | 100 | 1 | 1 | 0.025, 0.05, 0.1 |

